# Flexible multivariate linear mixed models for structured multiple traits

**DOI:** 10.1101/2020.03.27.012690

**Authors:** Hyeonju Kim, Gregory Farage, John T. Lovell, John K. Mckay, Thomas E. Juenger, Śaunak Sen

## Abstract

Many genetic studies collect structured multivariate traits containing rich information across traits. We present a flexible multivariate linear mixed model for quantitative trait loci mapping (FlxQTL) for multiple correlated traits that adjusts for genetic relatedness and that models information on multiple environments or multiple timepoints using trait covariates. FlxQTL handles genetic mapping of multivariate traits faster with greater flexibility compared to previous implementations.

---

Multivariate traits are increasingly common in genetic studies. A trait may be measured in multiple environments, at multiple timepoints (e.g. repeated measures), or under different treatments. Such a trait can be considered as a multivariate trait with spatial, temporal ^1, 2^ or more complex structures. For instance, body weight might be measured weekly, plant crop yield or biomass could be measured in multiple geographic locations for years, (in multi-environment trials, MET), and gene expression may be measured in different brain regions of the same rat. In humans it is common to study genotype-phenotype associations in population-based samples for genome-wide association studies (GWAS); in model organisms complex breeding designs are often used. Linear mixed models (LMMs) ^3–6^ are employed for controlling confounding due to genetic relatedness among individuals and are used to identify genetic loci contributing to quantitative traits of interest (QTL) ^7–13^. Multivariate LMMs (MLMMs) further enhance statistical power over univariate LMMs ^14^ because they can accumulate signals across traits ^8, 15^ with a common genetic locus.

While recognized as advantageous, fitting MLMMs is generally avoided because it is computationally challenging. Parameter estimation involves multidimensional optimization, there is a risk of reaching suboptimal solutions, and it is computationally expensive ^9, 14^. Existing algorithms (GCTA ^16, 17^, ASReml ^18^, WOMBAT ^19^, GEMMA ^14^) use the expectation-maximization (EM) method followed by second order schemes such as Newton-Rapson (NR), Average Information (AI), etc. to provide stable, fast convergence ^5, 14^. The computational complexity is *O*(*n*^3^*m*^3^) for EM and *O*(*n*^3^*m*^7^) for NR, AI (*m* traits, *n* individuals). This suggests that using GCTA, WOMBAT, and ASReml is not practical for GWAS with a large number of SNPs and a moderate number of individuals ^14^. GCTA can fit only up to 2 traits; ASReml and WOMBAT offer some flexibility in choosing some covariance structures and including modeling fixed effects of covariates. All have limitations in the number of traits they can handle and are much slower than GEMMA. None of these methods offer the ability to model spatial or temporal structure in the trait using trait covariates.

Here, we introduce FlxQTL, which can test associations between genetic markers and multiple correlated traits (**Supplementary Software** and https://github.com/senresearch/FlxQTL.jl). Our method is a MLMM that models the mean using a bilinear model of individual and trait co-variates (**Fig. 1**). The error term is the sum of a genetic random effect and a pure error component. The pure error term is correlated across traits but is independent across individuals. The genetic variance component is assumed to be proportional to the Kronecker product of a genetic kinship matrix and a trait kernel. In the case of MET, that can be interpreted as a random effect term comprised of the sum of many small random GxE effects (**Supplementary Note**). We assume

**Figure 1.**
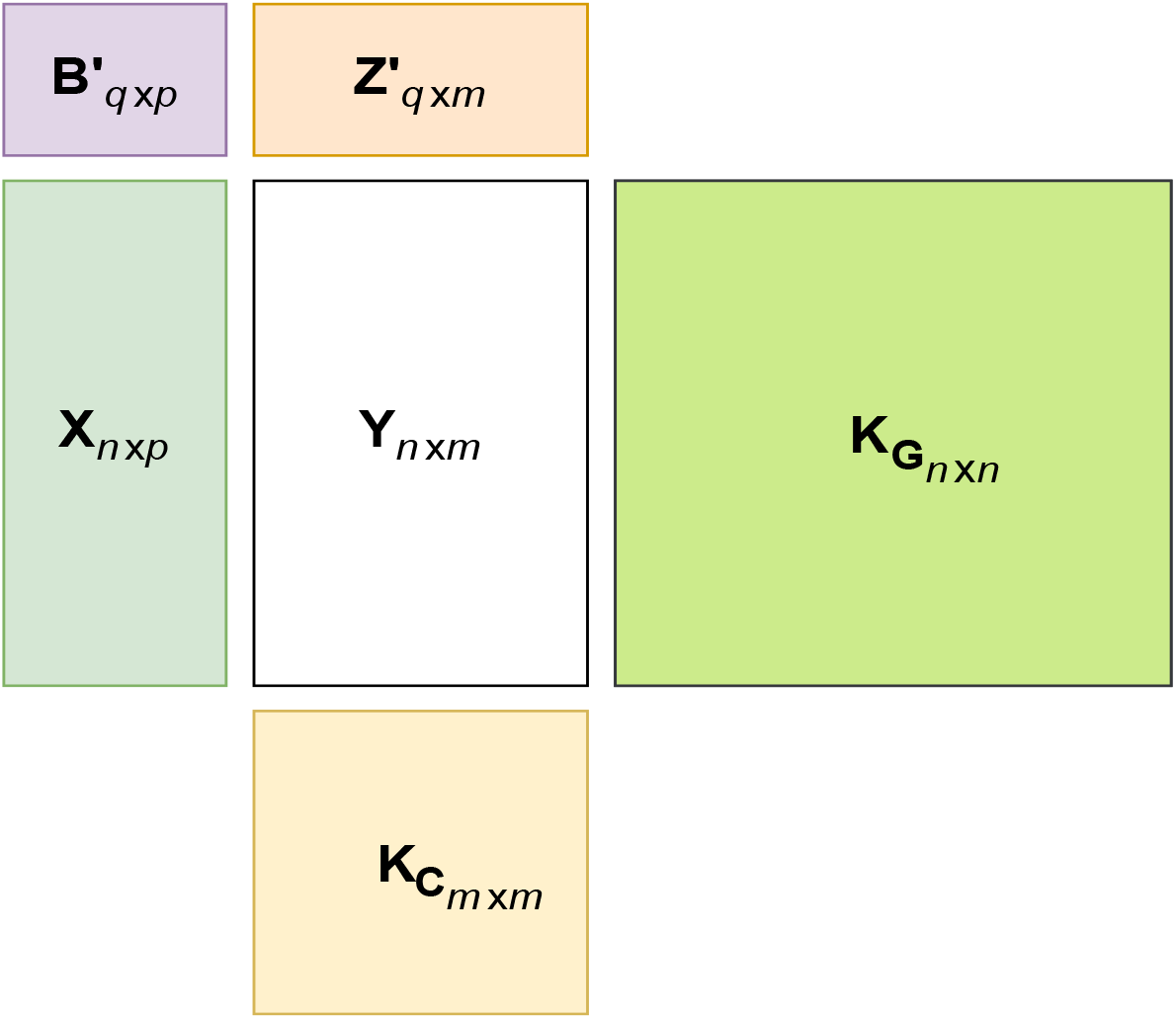
FlxQTL model components and their dimensions. The trait matrix (*Y*_*n*×*m*_) has information for *n* individuals (row) and *m* traits (column). In each dimension (by row and column), we have two paired components corresponding to fixed and random term respectively. The two components by row are a matrix of *p* genotypes of a marker (*X*_*n*×*p*_) to be tested and a kinship matrix (*K*_*G*_). The two components by column are a matrix of *q* trait covariates *Z*_*m*×*q*_, and a trait kernel *K*_*C*_. *B*_*p*×*q*_ is a matrix of fixed effects to be estimated. The two kernel matrices, *K*_*G*_ and *K*_*C*_ contribute to the variance component, and the two design matrices, *X* and *Z*, contribute to the mean components.

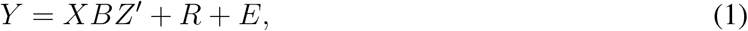

where *Y*_*n*×*m*_ is the response matrix for a trait measured in *n* individuals across *m* environments (or over *m* time points) or phenotypes of *n* individuals for *m* traits, *X*_*n*×*p*_ is a matrix of *p* genotype probabilities (or genotypes including the intercept) of a marker to be tested, and *Z*_*m*×*q*_ is a matrix of *q* trait covariates such as environment contrasts in MET, spline basis functions for modeling smooth temporal traits. *B* is a coeffcient matrix to be estimated, *R* is the matrix of genetic random effects, and *E* is the residual error matrix. The likelihood ratio test (LRT) to test the association between a marker and a multivariate trait of interest is expressed as a LOD score (− log_10_(*LRT*)). Genomewide thresholds for significance may be obtained from the null distribution of maximum LOD scores using a permutation scheme (Online Methods).

FlxQTL simplifies parameter estimation using its model structure (**Fig. 1**) and the choice of numerical methods. The trait covariates reduce the fixed effects matrix dimension from *p* × *m* for existing MLMMs ^8, 14^, to *p* × *q*, which is substantial when *m/q* is large. The trait kernel reduces a *m* × *m* random effects covariance matrix to a scalar parameter. With an unstructured pure error matrix, only 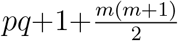 parameters are estimated, instead of *pm*+*m*(*m*+1) (*q* << *m*) for other MLMMs. FlxQTL has been tested with up to a 36-dimensional quantitative trait. Unlike NR and AI, a Speed restarting Nesterov’s accelerated gradient method requires no second derivative and provides stable linear convergence ^20^. We use an expectation-conditional maximization (ECM) ^21^ step, followed by the acceleration step using a tuned momentum coefficient, which behaves like the second derivative, to update parameters (**Supplementary Note**). These simplifications offer improved complexity compared to existing algorithms (Online Methods).

FlxQTL offers flexibility in handling varied trait structures and can analyze complex crosses if genotype probability data is available. The user can choose contrasts between trait covariates to influence how structured traits are analyzed. For example, GxE interactions can be analyzed using a contrast between sites in the *Z* matrix. One can use B-spline, wavelet, or Fourier basis functions in the *Z* matrix to model an environmental gradient or a time trend. By judiciously choosing the *Z* (trait covariate design) matrix, the user can model high-dimensional structured traits in MET, multiple related traits, and time-valued traits for a variety of model organisms as well as humans. Unlike some programs designed only for human genetics (maximum of three genotype states possible at a locus), if genotype probabilities are available, they can be used for QTL analysis in FlxQTL. This opens up the possibility of using 4-way crosses and multi-parental crosses such as heterogeneous stocks and the Collaborative Cross.

We compared the performance of FlxQTL with that of GEMMA on the Mouse HS1940 data distributed with GEMMA. The trait kernel and trait covariate matrices were set to be identity matrices in FlxQTL since they are not supported by GEMMA. We measured computation times for analyzing 3 traits and then increased the number of traits and the number of SNPs analyzed for both algorithms (Data processing and analysis in Online Methods). The difference in P values between the algorithms was negligible (**Fig. 2b** and **2c**). GEMMA was faster for a smaller number of traits (3 traits), whereas FlxQTL was 1.5-21 times faster than GEMMA when more traits were analyzed (6-12 traits) (**Fig. 2a**). These results show that FlxQTL offers fast implementation for high-dimensional traits.

**Figure 2.**
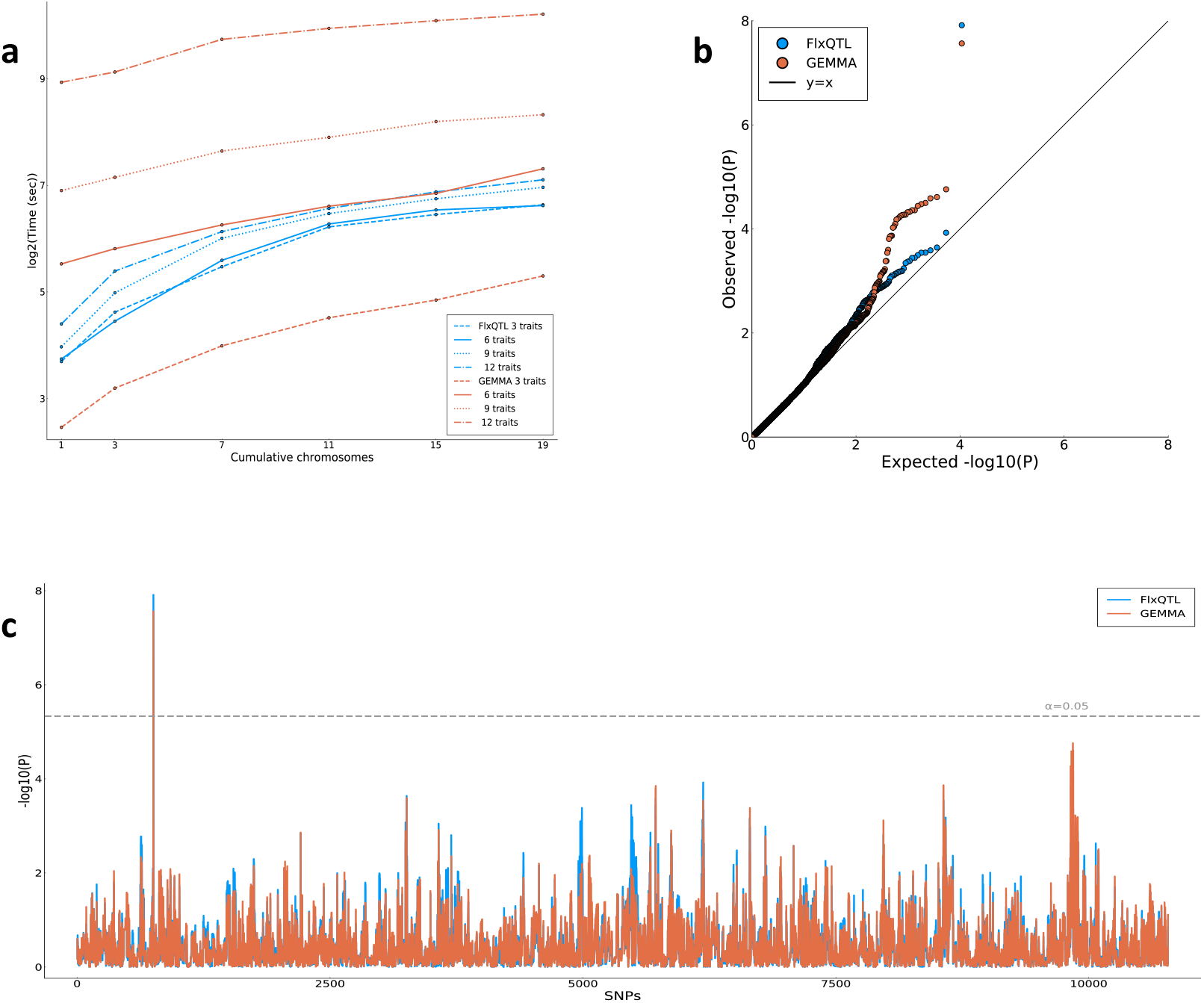
Comparison of FlxQTL and GEMMA using Mouse HS1940 data (n=1940 individuals, p=10783 SNPs) and simulated data. (**a**) Computation times in seconds on logarithmic scale (base 2) for parameter estimation as a function of the number of traits and SNPs. (**b**,**c**) Comparison of P values from the two algorithms using a quantile-quantile plot (**b**) and a genome scan plot (**c**) for 3 traits. The horizontal line is a Bonferroni adjusted threshold at the significance level of *α* = 0.05. Computing platforms: Linux Debian 4.19.37-5 (OS), Intel(R) Xeon(R) CPU E5-2630 2.4GHz (CPU), FlxQTL (Julia v.1.5.3 with 32 cores), GEMMA (v.0.98.3 with maximum 32 threads)

We evaluated the statistical performance of FlxQTL simulating 6-dimensional quantitative trait datasets using genotypes from *Arabidopsis thaliana* ^22^ studied in two sites for three years (Online Methods). We carried out two simulation studies on the contributions of a climatic relatedness matrix as a trait kernel and trait covariates, respectively. A comparison of power with, and without the climatic relatedness was made when including the same trait contrasts (**Supplementary Figs. 3-5**). We next compared power with and without trait covariates, allowing main effect QTL and QTL×site interactions, keeping those climatic relatedness matrices the same (**Supplementary Fig. 6**). The thresholds for the type I error rate were calibrated by the distribution of maximum LOD scores under the null model of only multivariate gene and environment random effects. The power was measured under the alternative model of existing large effects of QTL based on the thresholds. The overall result of assessing the trait kernel showed that including climatic relatedness as a trait kernel yielded only subtle differences in power. The result from the added value of trait covariates varying the strength of the genetic variance component demonstrated that in each case, the power was the highest when including trait covariates (**Supplementary Fig. 6**).

We applied FlxQTL on the data of recombinant inbred lines from *A. thaliana* by crossing populations, where the trait was the mean number of fruits per seedling planted in two distinct regions, Sweden and Italy, from July, 2009 to June, 2012. Due to the lack of formal statistical frame-works for QTL analysis in such reciprocal transplant experiments, Å gren et al. ^22^ performed six separate univariate QTL scans and dissected the effects post hoc. FlxQTL can directly test for the presence of genetic drivers of adaptation in replicated reciprocal transplants (the data processing and analysis is detailed in Online Methods, **Supplementary Tables 1** and **2** and **Supplementary Figs. 11-15**).

FlxQTL was applied to a more complex dataset, a four-way outbred mapping population in outcrossing switchgrass (*Panicum virgatum*). The trait was flowering time measured across 10 sites spanning a latitudinal gradient for 4 years (2016-2019). We considered it as a 36-dimensional multivariate trait since 4 site-year combination had missing data. These traits are highly correlated due to shared genetic and environmental contributions. We scanned for loci showing multivariate effects and picked the top three QTL. For those QTL, we performed an environment scan, for monthly averaged photoperiod in different sites and years. We found that the photoperiod influences one locus (QTL@6.6012 on Chromosome 5N) much more compared to the two other QTL (**Supplementary Fig. 16** and Data processing and analysis in Online Methods). This suggests one locus is more responsive to photoperiod and has implications for breeding.

As an example of a longitudinal trait, we analyzed body weight measured weekly from birth to 16 weeks in 1212 F_2_ intercrosses between Gough Island mice and WSB/EiJ ^2^. We used cubic splines with 4 degrees of freedom as the trait covariates. FlxQTL captured the weight trend and corresponding weight growth rate simultaneously and yielded the result similar to R/qtl ^23^. For two chromosomes, we performed two-dimensional genome scan for further investigation and found several more QTL. (Online Methods and **Supplementary Figs. 17-21**).

Our results demonstrate that the model and estimation algorithm are effective in analyzing higher dimensional structured traits with fast parameter estimation. On the simulation study, FlxQTL increases power making use of information from trait covariates. Our method appears to be insensitive to a choice of trait kernel matrices in our studies so far, and this may be worth further investigation.

The underlying trait model assumes a joint multivariate normal distribution and is not ideal for traits such as count data with lots of zeros, which do not fit this scenario. Some traits may need transformation to better fit the multivariate normal assumption. Our method assumes that the multivariate trait has no missing data and that genotypes are either completely observed or imputed, or genotype probabilities are used. If phenotype data contain missing values, one can use imputation ^24^. With the growth of more complex and high-throughput phenotyping, we expect the use of our models to help shed light on the genetic control of structured multivariate traits.

## METHODS

Methods and any associated references are available in the Online Methods.

## ONLINE METHODS

### Code availability

A Julia software implementation of FlxQTL is available at https://github.com/senresearch/FlxQTL.jl.

### FlxQTL model

Let *Y*_*n*×*m*_ be a trait measured in *n* individuals across *m* environments (or over *m* time points), or phenotypes of *n* individuals for *m* correlated traits. *X*_*n*×*p*_ is a matrix of *p* genotypes including the intercept (or genotype probabilities) of a marker to be tested and can also optionally include individual level covariates such as sex for animals or cytoplasm origin for plants. *Z*_*m*×*q*_ is a matrix of *q* trait covariates such as (site) contrasts, (orthonormal) basis functions, etc. Our model for the trait is:

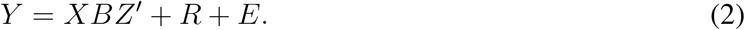

The two independent random effects follow matrix variate normal distributions:

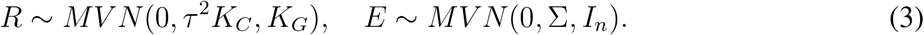

One can easily see that compared to standard multivariate regression, where *Z* = *I*_*m*_, *q* = *m*, our model reduces the size of *B* from *p*×*m* to *p*×*q*. This dimension reduction is consequential when *q* ≪ *m*, as is often the case when working with higher dimensional trait data.

In the first variance component, *K*_*G*_ is a genetic relatedness (kinship) matrix between individuals computed by background genetic markers, and *K*_*C*_ is a trait kernel generated by extra information associated with the traits; in MET, the trait kernel can be a relatedness matrix derived from high-dimensional environment (or site) information such as daily minimum, or maximum temperature, precipitation, etc. We show that for MET, the variance component, *R*^*v*^, composed of many infinitesimal *G*×*E* effects, has variance proportional to *K*_*C*_ ⊗ *K*_*G*_ (**Supplementary Note**). *τ* ^2^ is a scalar parameter for reducing the dimension of a covariance matrix, which is unknown in other studies, by assuming it is proportional to *K*_*C*_. If no information is available on the multiple traits, then *K*_*C*_ = *I* is advisable.

The second variance component assumes correlations among traits but independent and identically distributed (iid) noise between individuals. The common error covariance matrix is Σ which assumes to be unstructured (no constraints). The two kernel matrices are precomputed from *a priori* information, so that only *τ* ^2^ and Σ are estimated. The MLMM in FlxQTL is then modeled (in vectorized form) by a multivariate normal distribution whose covariance matrix is the sum of two Kronecker products for the two independent random effects:

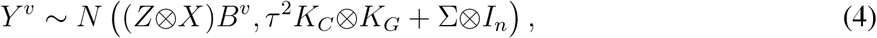

where the superscript *v* denotes the vectorization of a matrix. LOD scores (− log_10_(*LRT*)) are calculated by log-likelihood difference between the full model and null. We use permutations (at least 1000) to calculate empirical genomewide LOD thresholds as follows: 1. The traits are rotated by the eigenvectors of *K*_*G*_ (and *K*_*C*_(≠ *I*)). 2. The rotated traits are divided by the square root of their variance-covariance matrix to make them iid. 3. They are shuffled by row and transformed back by multiplying by the square root of the covariance matrix. 4. A genome scan is performed on this shuffled trait, and the maximum LOD for each permutation is stored. Note that LOD scores using trait covariates (*Z*) are lower than those implemented by conventional MLMMs (i.e. *Z* = *I*), but the genomewide significance threshold is also correspondingly lower.

#### Computational costs

The eigen decomposition of the kernel matrices is done only once; this has computational complexity *O*(*n*^3^ + *m*^3^) ≈ *O*(*n*^3^) (*m* << *n*), the same as existing MLMMs. A LOCO (Leave One Chromosome Out) scheme ^7, 9, 25^ has complexity *O*(*dn*^3^) ≈ *O*(*n*^3^) (*d* ≪ *n*) for *d* chromosomes and does not change complexity appreciably. In practice, genome scans with the LOCO scheme take longer depending on the data (about 30 % longer) than that without LOCO. For *t*_0_ ≫ *t*, where *t*_0_, *t* are the maximal number of iterations of ECM and ECM embedded in the Speed restarting Nesterov’s scheme, respectively, the computational complexity of two step implementation, which runs ECM for finding ‘good’ initial values followed by ECM embedded in the Speed restarting Nesterov’s scheme, is *O*((*c*_1_*qn*^2^ +(*m*^3^ +*c*_1_*m*^2^)*n*)(*t*_0_ +*t*)) per marker for *n* individuals, *c*_1_ covariates. The whole computational complexity including the eigen decomposition and rotation by corresponding eigen vectors is now *O*(*n*^3^ + (*m* + *c*_1_)*n*^2^ + *s*(*c*_1_*qn*^2^ + (*m*^3^ + *c*_1_*m*^2^)*n*)(*t*_0_ + *t*)) with *s* markers.

#### Julia implementation

FlxQTL is implemented in Julia ^26^, a relatively new programming language, well-suited for developing and implementing algorithms for large datasets. Julia has a just-in-time (JIT) compiler, which allowed us to prototype algorithms with an ease comparable to scripted languages like R, and with a speed approaching compiled languages. Julia has native support for distributed and parallel computing, and users can take advantage of multiple cores or a high-performance computing (HPC) system to achieve a significant speedup. Genome scan with millions of markers involving moderately high-dimensional traits is possible.

### Data processing and analysis

#### Mouse HS1940 data

SNPs were filtered out by minor allele frequency (MAF) less than 2% since we found GEMMA automatically processed the data with that criterion. For a fair comparison, we used a centered genetic relatedness matrix computed by GEMMA, but it needed some adjustment by adding very small value (0.00001 * *I*) to make sure the matrix should be positive definite. The trait data consisted of 3 traits with missing values and was imputed using R package, mice, with a ‘pmm’ option (predicted mean method). To compare the computation time between FlxQTL and GEMMA as a function of the number of traits and SNPs, we created new trait (triplets) by shuffling the trait data by individual preserving the correlation among three traits and added them to the existing traits. This produced 6, 9, and 12 simulated traits. We also generated 19 cumulative genotype datasets from the mouse data, that is, starting from a chromosome 1 SNP dataset (950 SNPs), building up a new dataset with 4-chromosome increments and reaching the whole SNP dataset (10783 SNPs). The median time for the genome scan without LOCO was obtained from 64 runs on each simulated trait data.

#### Gough Island mouse data

This is a F_2_ intercross between Gough Island mice and WSB/EiJ. We computed genotype probabilities, excluding sex chromosomes, from R/qtl adding pseudo markers in every 1cM between two markers. Traits were imputed using R-mice package with the ‘pmm’ option and then were standardized by overall mean and standard deviation. We employed a 4 degrees-of-freedom cubic spline from R-splines package to produce trait covariates (*Z*) to capture the trend of weekly body weights and corresponding weight growth rate simultaneously. Here, we set the trait kernel to be an identity matrix. A genetic relatedness matrix was computed from genotype probabilities using a linear kernel. The thresholds at *α* = 0.1, 0.05 were estimated from 1000 permutations. The results from the one-dimensional multivariate genome scan by FlxQTL was largely consistent with that ^2^ from R/qtl using univariate linear regression, except for Chromosome 6 and 7 perhaps due to polygenic effects. Since one QTL in Chromosome 7 was known to be crucial, we implemented a two-dimensional genome scan for Chromosome 7, as well as 8 and 10 for comparison. FlxQTL detected several more QTL than R/qtl without requiring further analysis such as multiple-QTL analysis.

#### Arabidopsis thaliana data

We employed R/qtl to add pseudo markers in every 1cM between two markers and dropped one of two consecutive markers when they are identical. The trait data were imputed using the ‘pmm’ option in R-mice package and were standardized by column. A total of 699 markers across 5 chromosomes were genotyped for 400 recombinant inbred lines, and a fitness trait was measured at two sites (Italy and Sweden) for three years (6-dimensional multivariate trait). The trait covariate matrix (*Z*) set to be a matrix with an intercept column, and a contrast between sites (1’s for Sweden and -1’s for Italy). We computed both a genetic and a climatic relatedness matrix using Manhattan distance. The climatic relatedness matrix was computed using daily range soil or air temperature, precipitation, etc. However, the difference between with and without climatic relatedness matrix appears to be small in our simulation studies (**Supplementary Fig. 7**). We performed 1-, 2-dimensional QTL analyses with the LOCO option followed by multiple QTL analysis, i.e. a stepwise model selection approach by forward selection and backward elimination adding or dropping one or two QTL for each scan, respectively. The 95% cutoff was estimated by permutation test and was used for a penalty term in the stepwise model selection since the false positive rate is maintained at the rate of *α* in the case of no QTL and with the search restricted to models with no more than one QTL ^27^. Our result revealed one more significant QTL in three chromosomes each but one less in two chromosomes each, by and large, agreeing with the existing result, with improved interpretation and without requiring secondary analysis of QTL across chromosomes ^22^.

#### Switchgrass (Panicum virgatum) data

Genotype probabilities were computed in every 1cM between two markers via R/qtl. Traits and climate information data had no missing values, so that the whole dataset comprises 6118 genetic markers by 750 individuals for 36 quantitative traits of flowering time as a combination of 10 latitudinal sites and 4 years from 2016 to 2019, as well as a matrix of 12 by 36-monthly photoperiod. Three significant QTL (two in Chromosome 5N and one in Chromosome 4K) were selected by 1D-genome scan with the LOCO option. For each QTL, an environment scan was performed by regressing each monthly photoperiod factor under the alternative of existing an environmental factor that affects a QTL after the null scan. For computational efficiency, traits and monthly photoperiod data were centered by column means and scaled by overall standard deviations of the centered data.

### Simulation study

We simulated data with the same overall structure and features of the Arabidopsis dataset ^22^. A 6-dimensional multivariate trait was simulated with random GxE effects based on genotype data and daily soil minimum and maximum temperature data from those sites and years. Soil daily range temperatures (the difference between maximum and minimum temperatures) were rearranged into a 3 year by 2 location combination, i.e. 365 days by 6 environments, and were standardized by overall mean and standard deviation. To generate null trait datasets, we selected at random about 3.5% of the genetic markers (about 25 markers of total 699 markers) and 8.2% of 365 soil daily range temperatures (about 30 days) and gave their interaction effect sizes drawn from a normal distribution with zero mean and variance *τ* ^2^ that varies from small to large (**Supplementary Note**). An iid noise indicating random error was added to the null trait data; the covariance matrix of the error term was varied. Both random and error terms were respectively standardized. To generate a fixed effect QTL to the null data, we sampled one genetic marker not selected among the random effects above and multiplied it by fixed effect sizes varying from small to large and the orthonormal site contrast matrix (*Z*). For simplicity, we only considered one fixed effect QTL. We carried out 1000 simulations under the null to attain a threshold at *α* = 0.05 from the distribution of maximum LOD scores and measured power under the alternative for each instance given the trait kernel. The first simulation study to assess power for the effect of climatic relatedness varied with values of *τ* ^2^ for each fixed effect size matrix (*B*), which in total resulted in 100,000 experiments. We then narrowed down to the feasible range of fixed effect sizes by excluding the effect sizes producing zeros and ones of power. Within that range, simulations were performed to establish a comparison between inclusion and exclusion of the climatic relatedness matrix (*K*_*C*_) changing *τ* ^2^. Since corresponding results showed slight distinctions in power, we tackled unusual scenarios in genome scan, where *K*_*C*_’s are an autocorrelated matrix, an extreme case of positive definite matrix, etc. The results revealed analogous power to those shown in prior experiments. The second simulation study to assess the effect of trait covariates (*Z*) was implemented in the similar manner. In the given range of fixed effect sizes, we compared three cases: non-identity contrasts (site contrasts in FlxQTL), an identity contrast (*Z* = *I* in FlxQTL), Julia version MLMM. Note that the Julia version of MLMM was developed as an alternative of GEMMA for ease of comparison with our method.

## Acknowledgements

This work was funded by NIH grants R01GM123489 (S.S., H.K.) R01AI121144, and P30DA044223 (S.S.); Department of Energy award DESC0014156, IOS0922457 and IOS1444533 (T.E.J.); NSF Awards DEB1022196 and DEB1556262 (J.K.M.). The work conducted by J.L. was funded by the US Department of Energy Joint Genome Institute, which is supported by the Office of Science of the US Department of Energy under Contract No DE-AC02-05CH11231. We thank Jon Å gren and Grey Monroe for providing climate data from the *Arabidopsis thaliana* reciprocal transplant experiments; K. Broman and B. Payseur for the Gough Island mouse data.

## Author Contributions

S.S. conceived the idea; H.K. and S.S. designed the study. H.K. wrote the first draft of the paper, implemented the algorithms and analyzed the data. G.F. contributed to developing the software, J.K.M., T.E.J. and J.L. contributed data and interpreted the results, and all authors revised the manuscript.

## 1 Supplementary Figures

**Figure 3.**
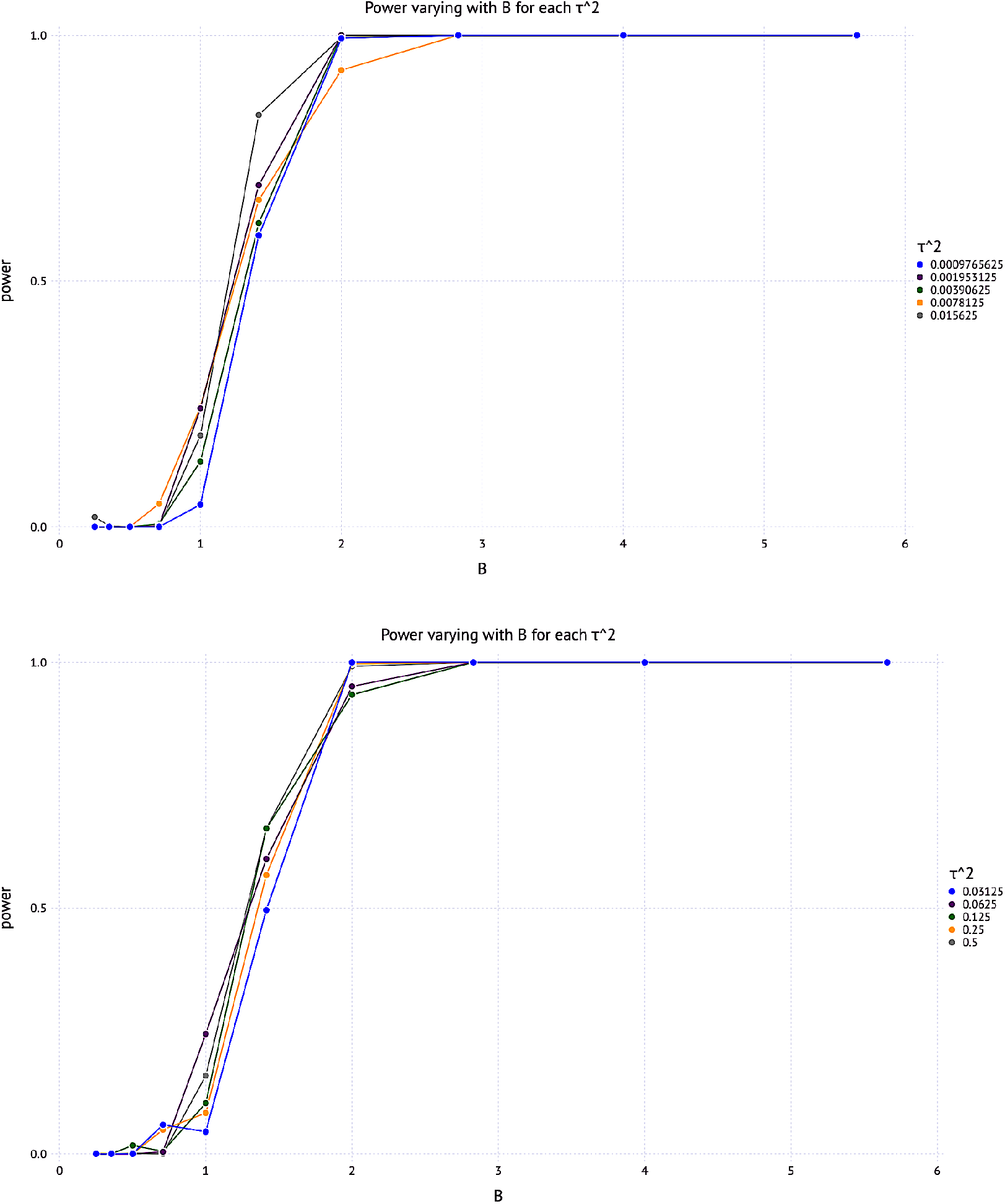
Power varied by *τ* ^2^ on the simulated data from Arabidopsis thaliana. The climatic relatedness matrix (*K*_*C*_) as a trait kernel was computed by 3 year by 2 site soil range temperature data. True effect size matrices *B*’s varied by the following formula: 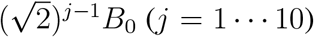 for *B*_0_=[0.25 -0.25;0.25 0.25], randomly selected from {±0.25}.

**Figure 4.**
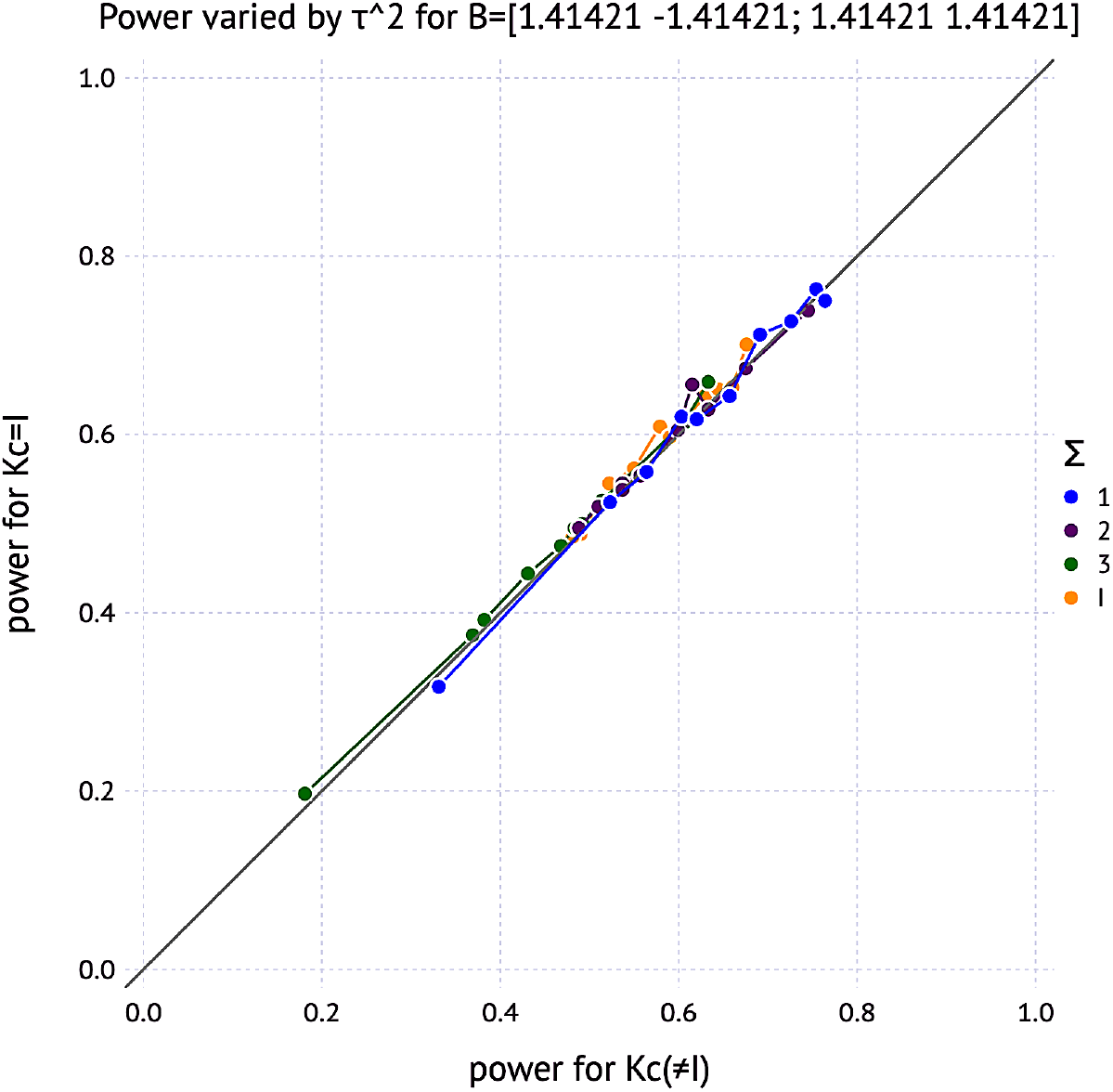
Power comparison of a non-identity *K*_*C*_ and an identity *K*_*C*_(= *I*) on the simulated data from Arabidopsis thaliana depending on unstructured environment errors (Σ). *K*_*C*_ was computed by 3 year by 2 site soil daily range temperature data. 6 × 6 matrices of Σ_*i*_ for *i* = 1, 2, 3 are selected using following 3 × 3 sub-matrices: *A, C* (1’s on the diagonal, *a, c* on off-diagonal entries, respectively), *B* (*b* in all entries). Σ_1_ = [A B;B A], where *a* = 0.4, *b* = 0.07, Σ_2_ =[A B;B A], where *a* = 0.3, *b* = 0.07, Σ_3_=[A B;B C], where *a* = 0.3, *b* = 0.0, *c* = 0.1. *I* is an identity matrix. The result shows subtle differences in power depending on Σ’s

**Figure 5.**
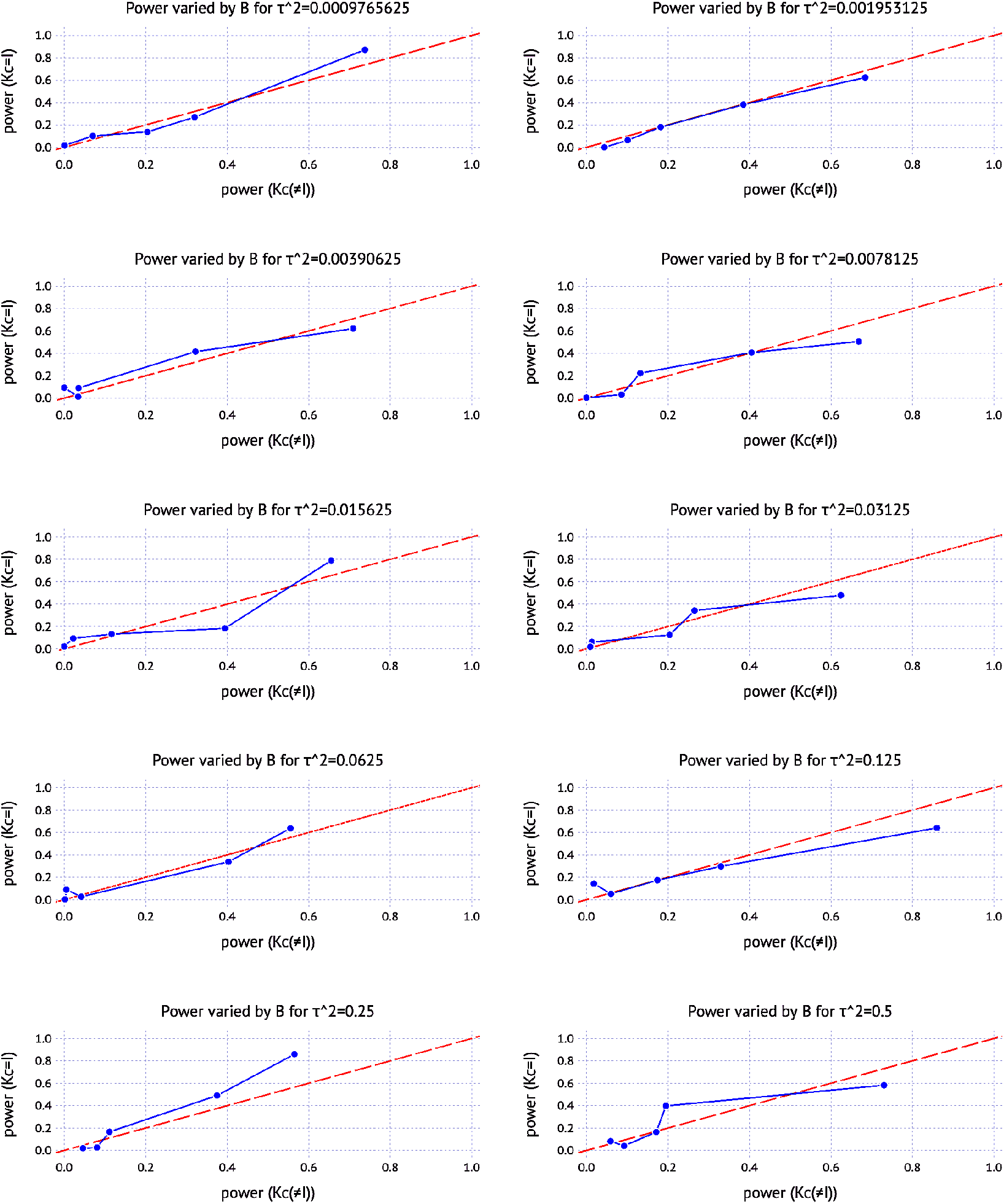
Comparison of power between a non-identity *K*_*C*_(≠ *I*) and an identity *K*_*C*_(=*I*) on the simulated data from Arabidopsis thaliana. The non-identity *K*_*C*_ was computed by 3 year by 2 site soil daily range temperature data.

**Figure 6.**
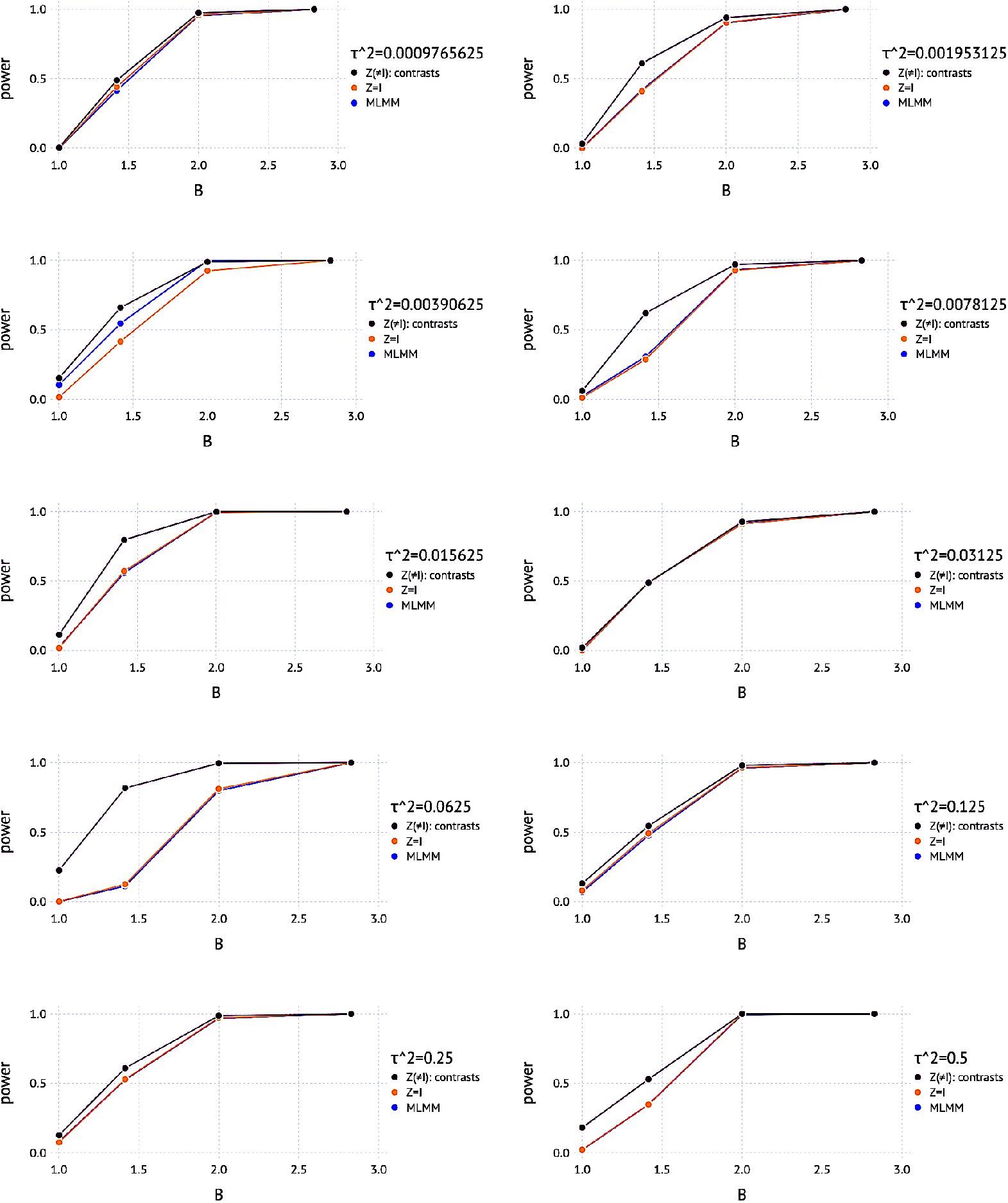
Contribution of trait covariates (*Z*) to power on the simulated data from the Arabidopsis thaliana. *K*_*C*_ was computed by 3 year by 2 site soil daily range temperature data. 1. *Z*(≠ *I*): a 6 × 2 matrix of contrasts including the intercept and the site contrast of −1’s and 1’s for Italy and Sweden, respectively. 2. *Z* = *I*: a 6 × 6 identity matrix. Both were run in FlxQTL. 3. MLMM: the julia-version MLMM as an alternative for GEMMA.

**Figure 7.**
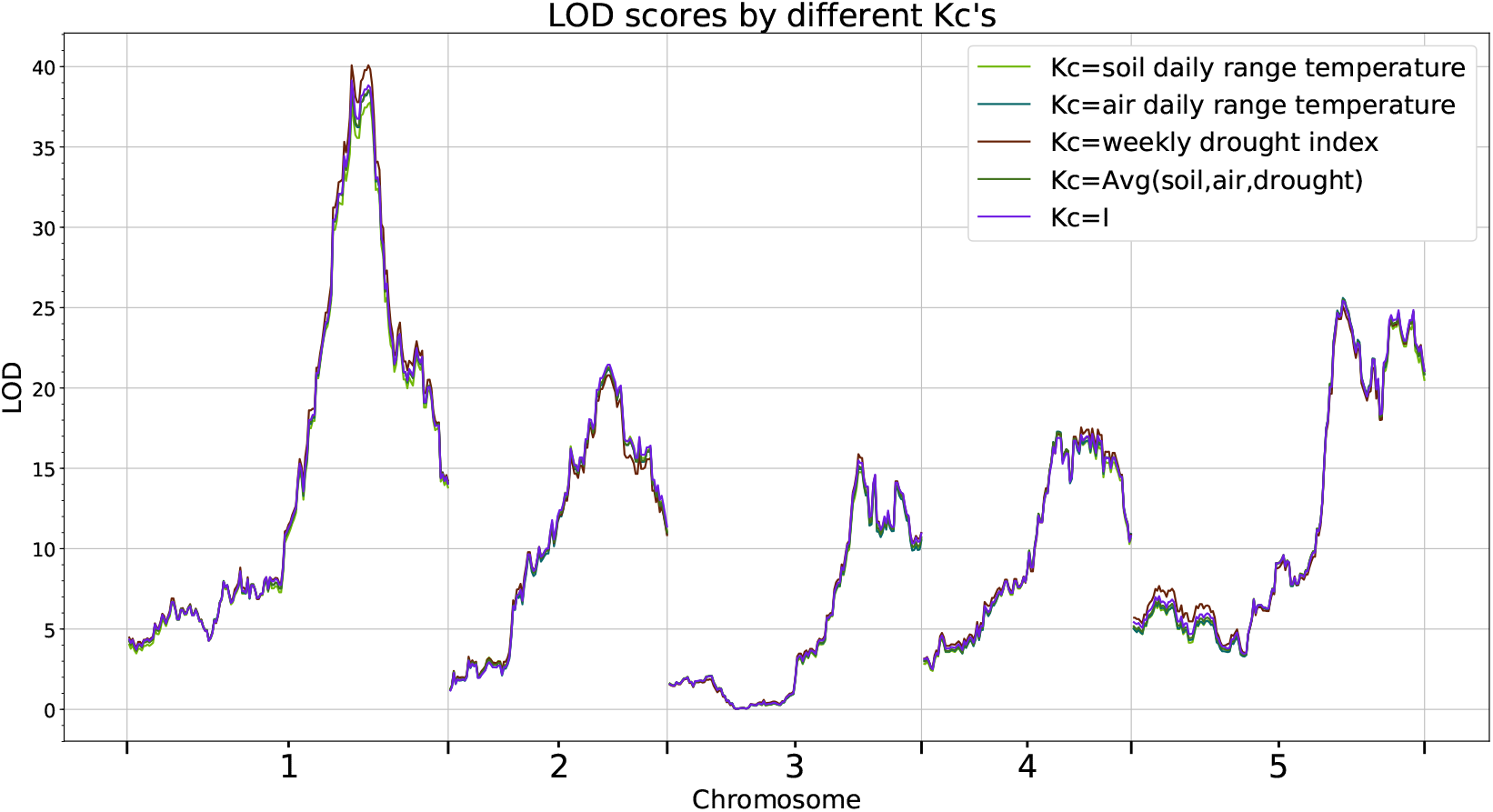
Multivariate genome scan for Arabidopsis thaliana: Comparison of LOD scores depending on various climatic relatedness matrices (*K*_*C*_). The trait was fitness (mean number of fruits per seedling planted) considered as 6 sites (a 3 year by 2 site combination). *K*_*C*_’s were respectively generated by using soil/air daily range temperatures, weekly drought indices in two sites (Sweden and Italy) for 3 years. The trait covariates (*Z*) were contrasts containing 1’s for overall mean for 6 sites and -1’s (Italy), 1’s (Sweden) for mean difference between sites to measure GxE interactions. The result appears to be insensitive to the choice of *K*_*C*_’s.

**Figure 8.**
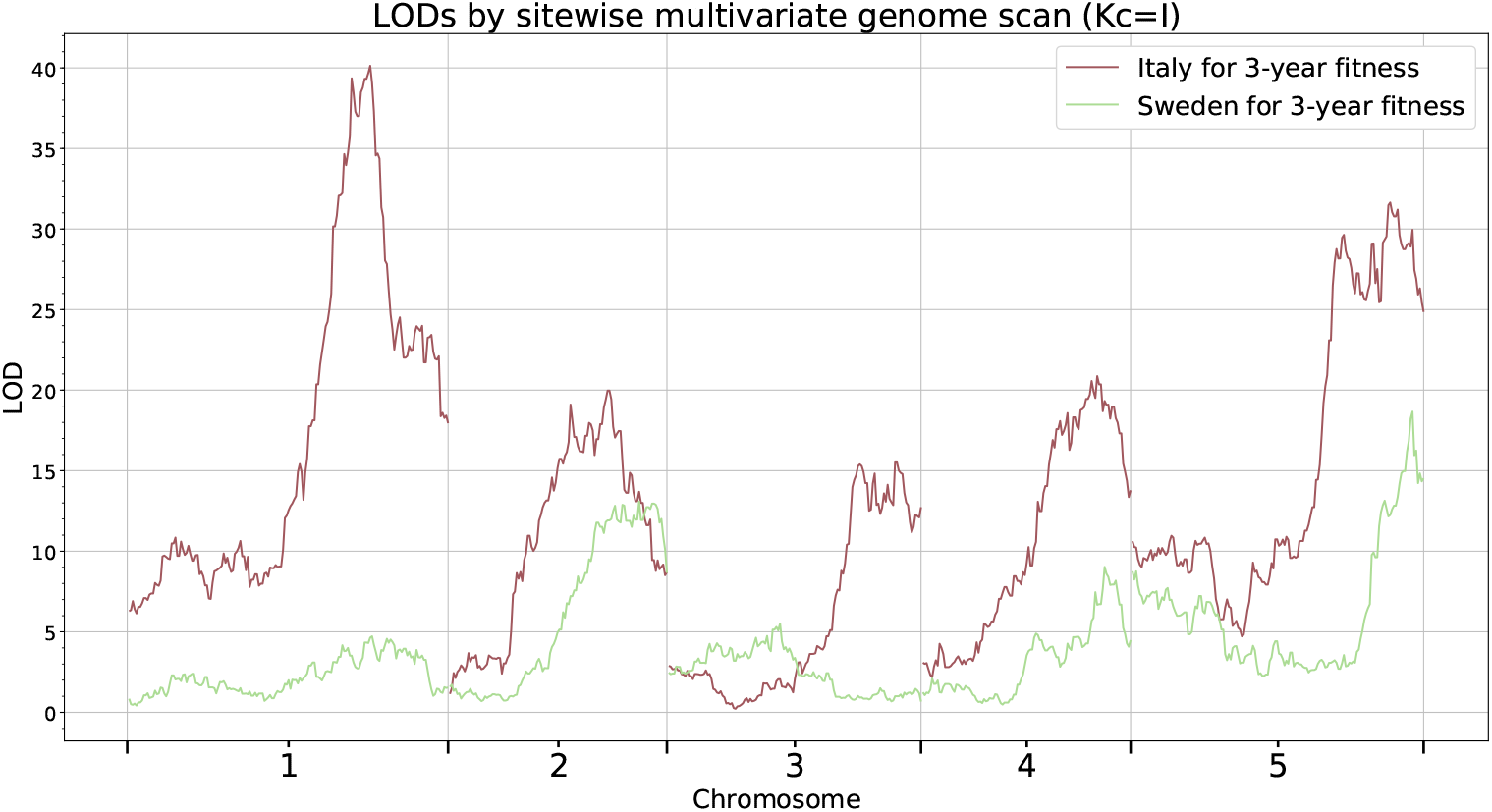
Site-wise multivariate genome scan for Arabidopsis thaliana: Comparison of LOD scores for Italy and Sweden, where *K*_*C*_ = *I* due to the result in Supplementary Figure 7.

**Figure 9.**
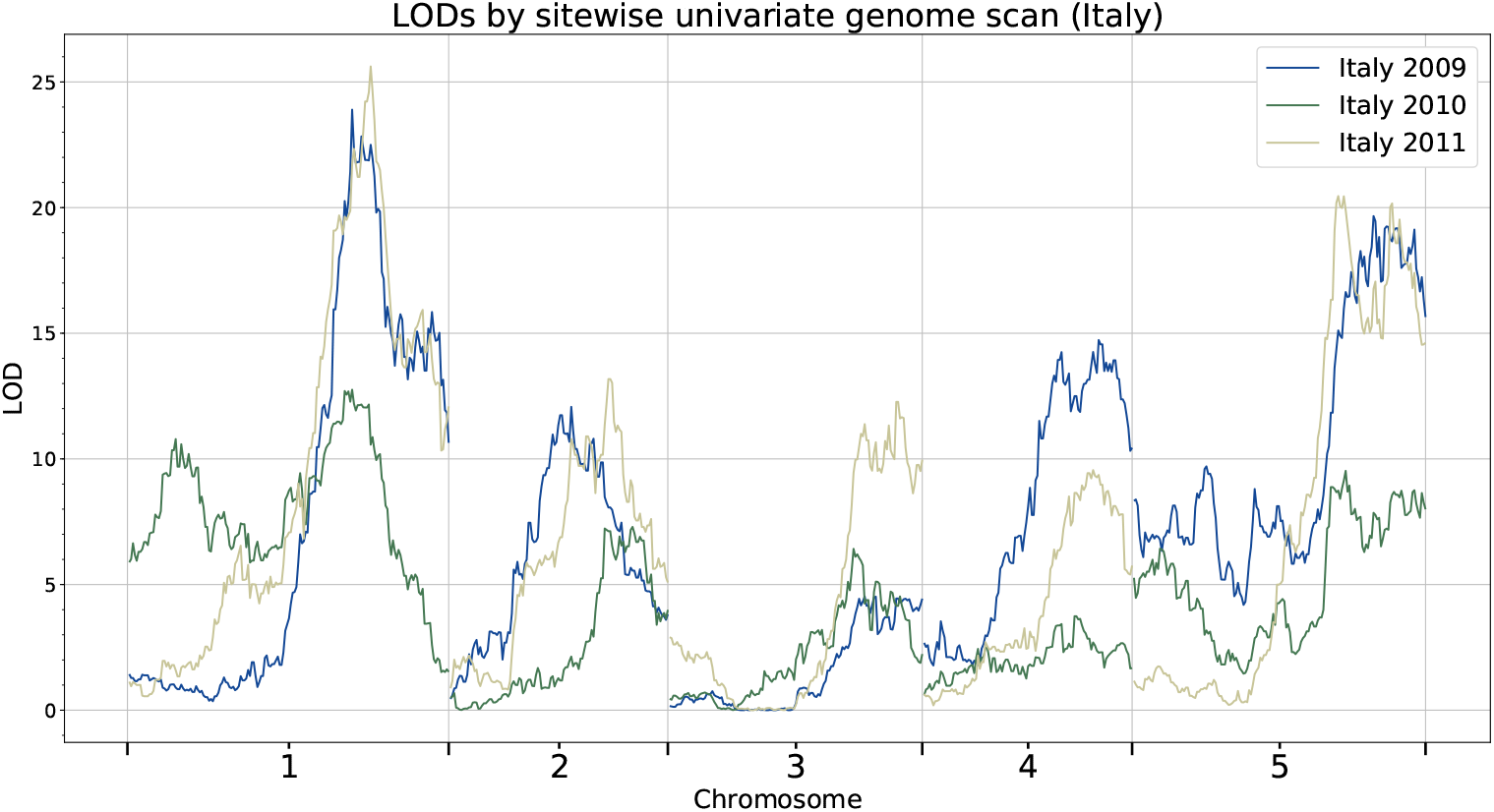
Univariate genome scan for Arabidopsis thaliana in Italy from 2009 to 2011.

**Figure 10.**
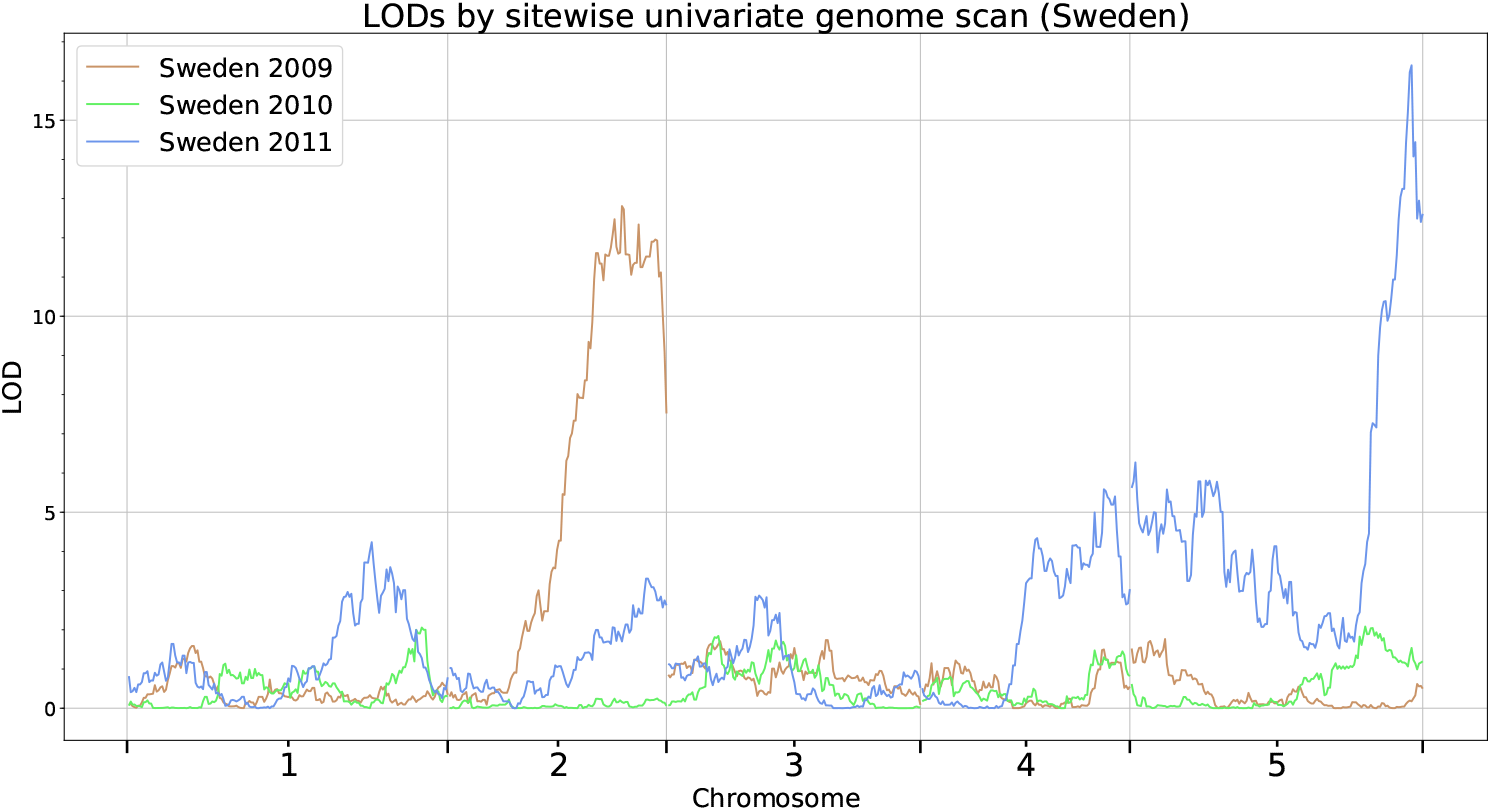
Univariate genome scan for Arabidopsis thaliana in Sweden from 2009 to 2011.

**Figure 11.**
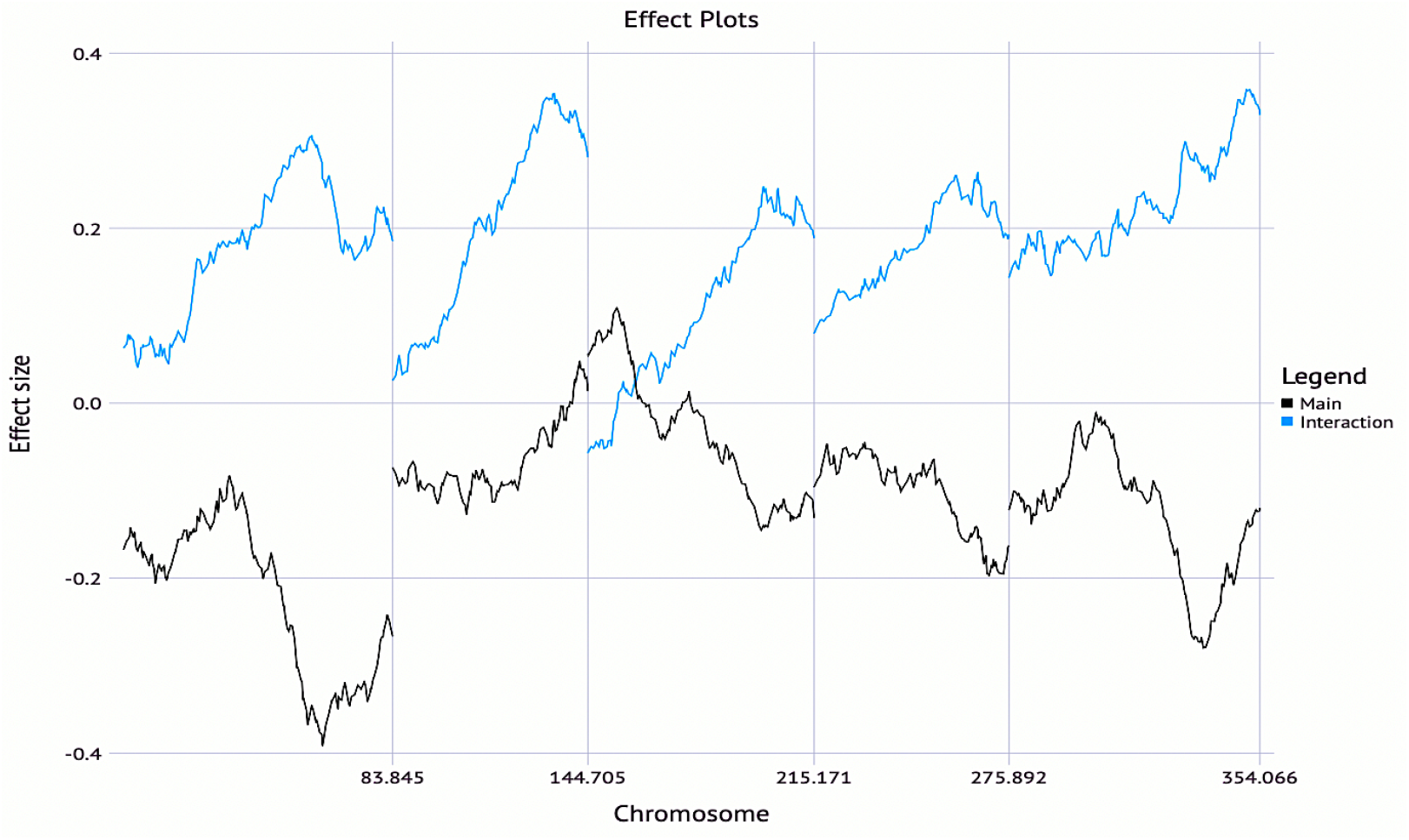
Effect plots for Arabidopsis thaliana with *K*_*C*_ = *I* for 6 quantitative traits as described in Supplementary Figure 7. Negative values in most main effects imply that the Swedish allele, on average, underperforms. Positive values in most interaction effects indicates home allele advantage; that is, the Swedish genotype in Sweden performs better than that in Italy.

**Figure 12.**
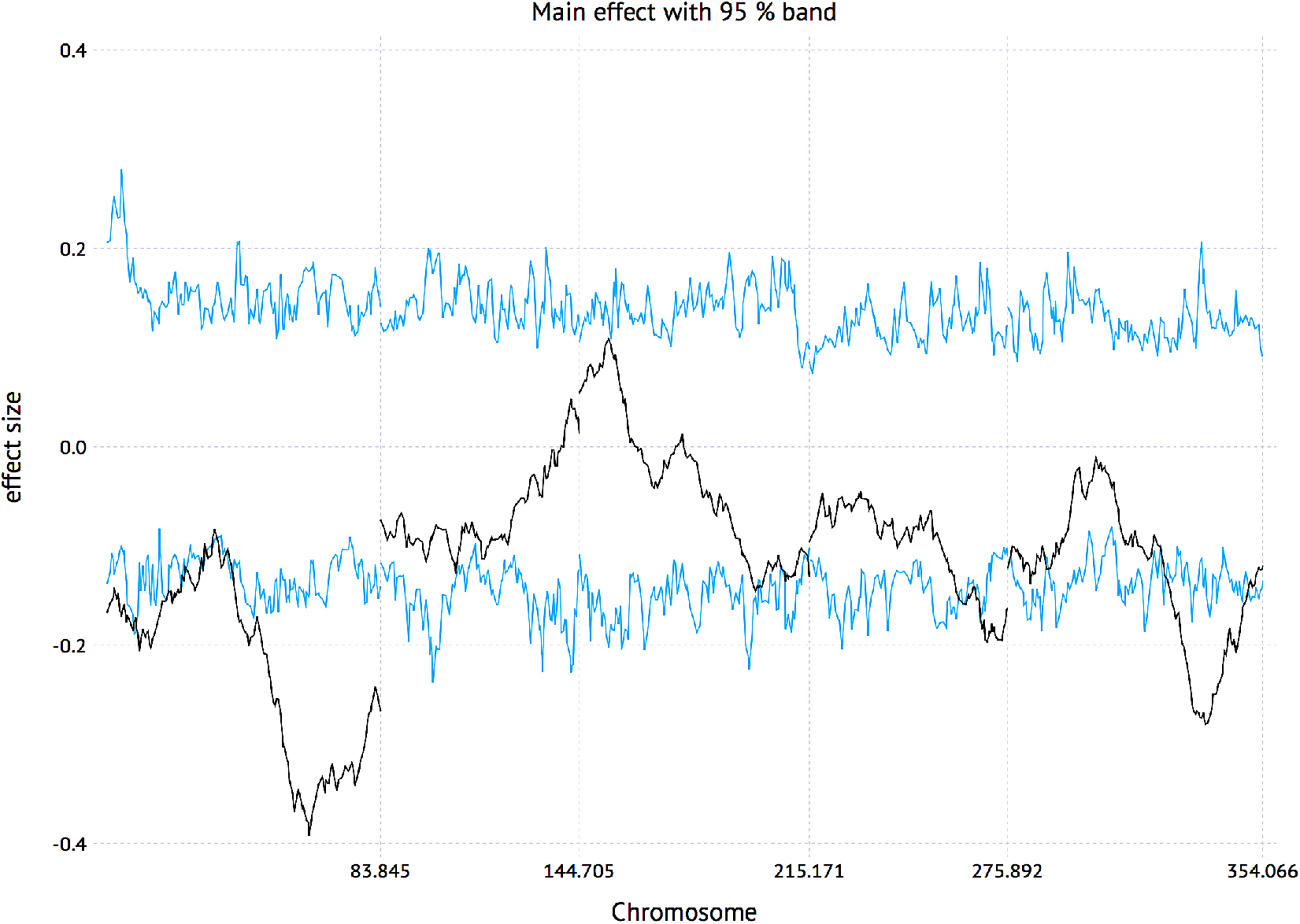
Main effects with 95% band for Arabidopsis thaliana. The band was obtained by 100 permutations

**Figure 13.**
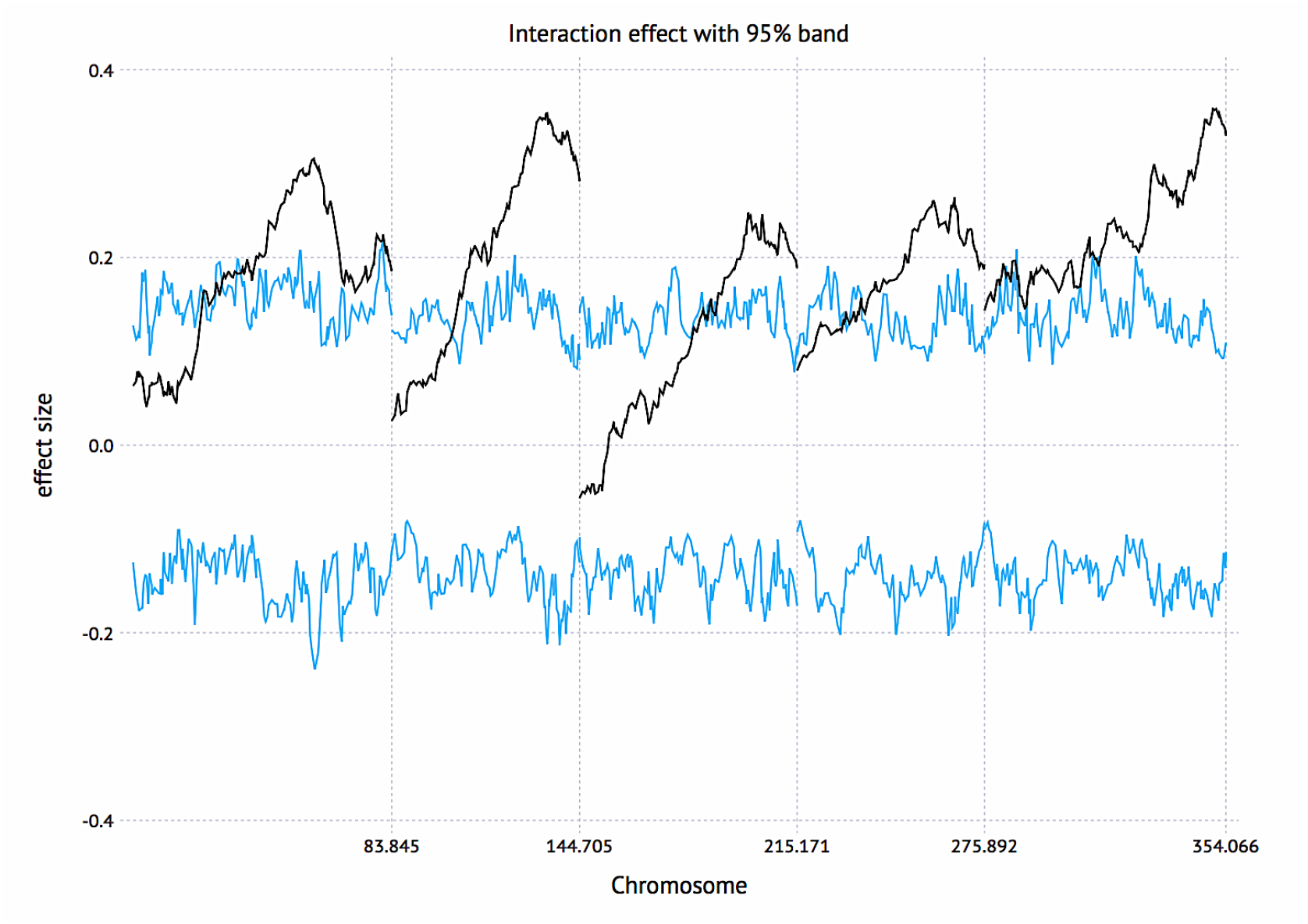
Interaction effects with 95% band for Arabidopsis thaliana. The band was obtained by 100 permutations

**Figure 14.**
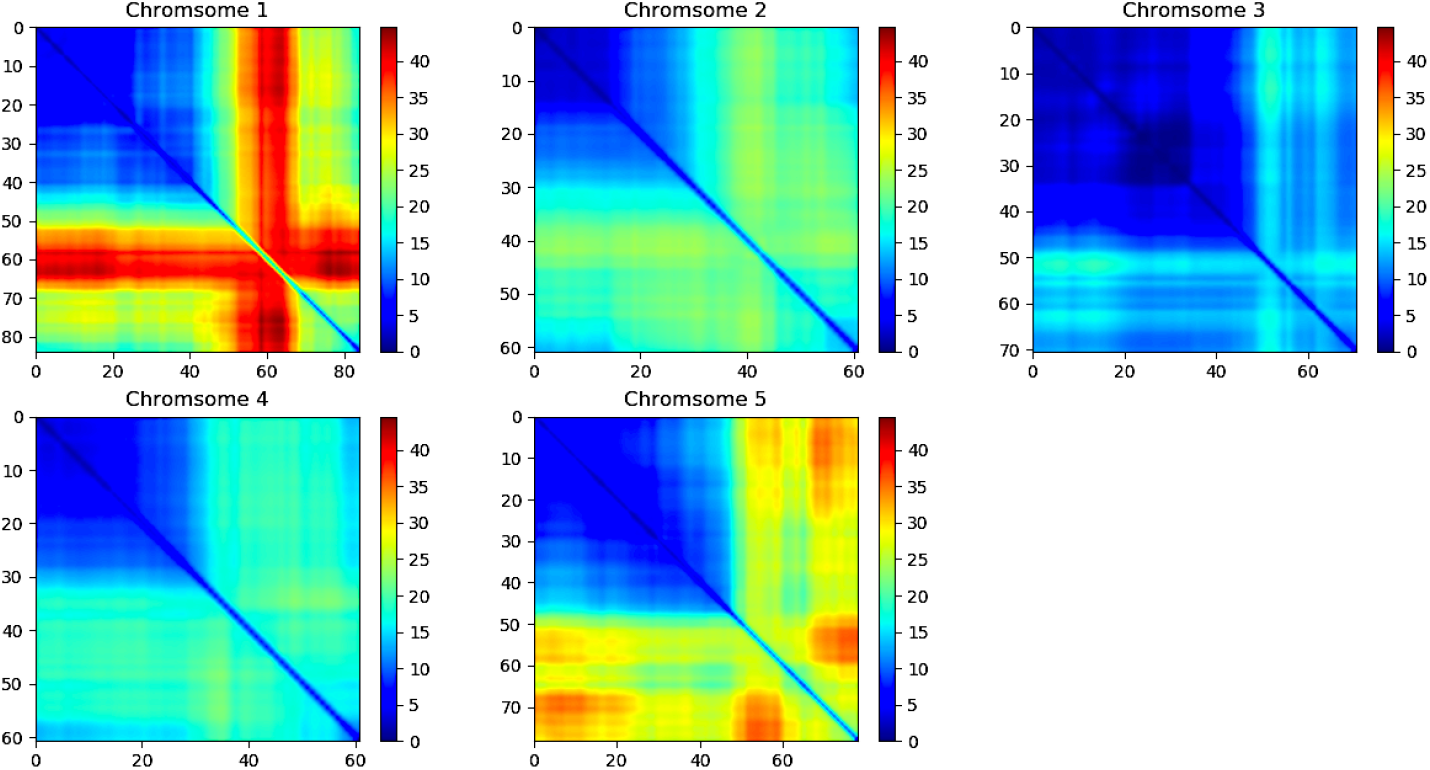
2-dimensional (2D) multivariate genome scan for Arabidopsis thaliana.

**Figure 15.**
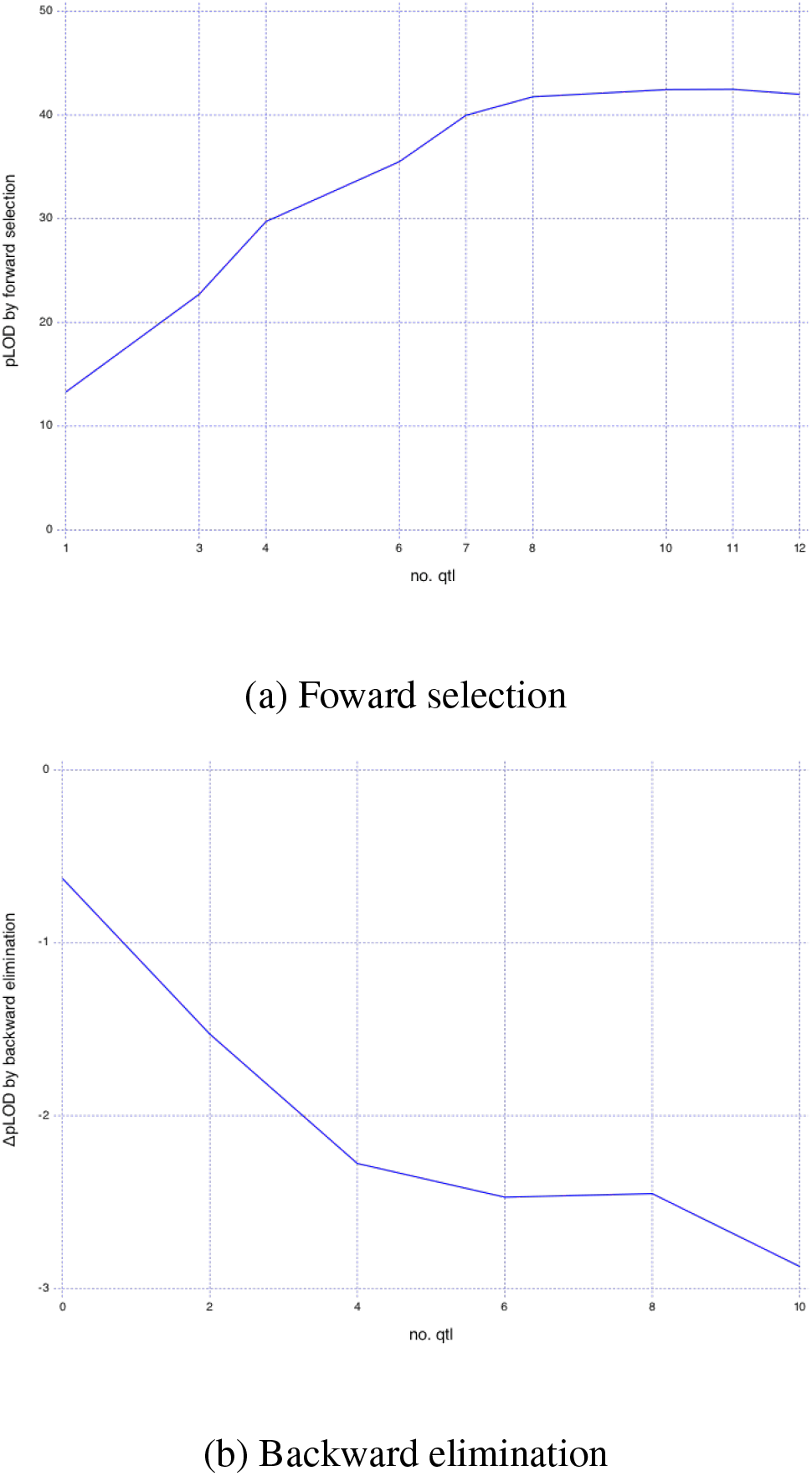
QTL selection for Arabidopsis thaliana (multiple QTL analysis). A model selection approach by forward selection and backward elimination was performed to finalize significant QTL from candidate QTL selected in Supplementary Figure 7, 14. A selection criterion is the penalized LOD score for additive models equivalent to BIC ^27^.

**Figure 16.**
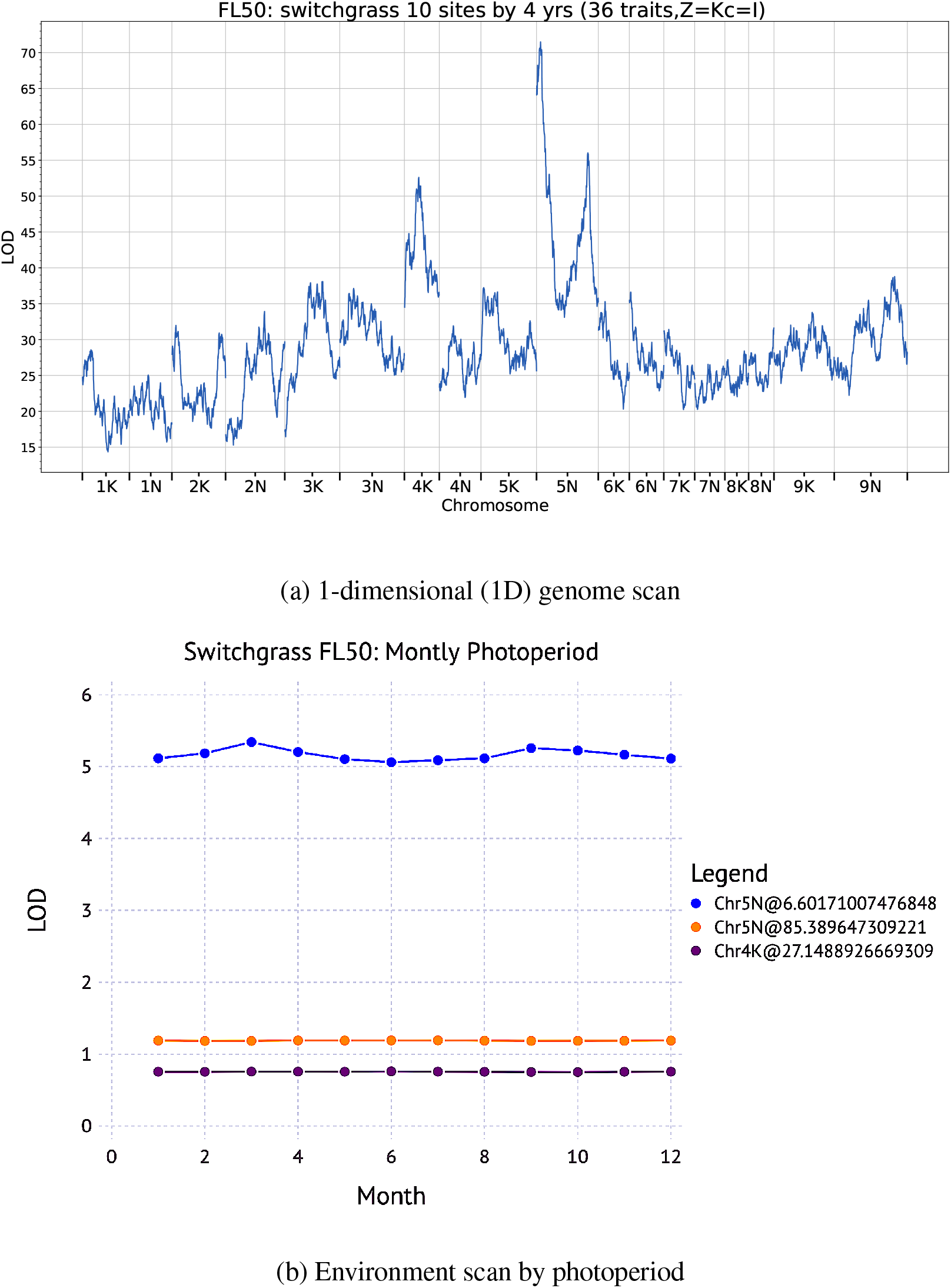
Switchgrass FL50. The environment scan (b) was done for the top 3 QTL selected from the genome scan (a)

**Figure 17.**
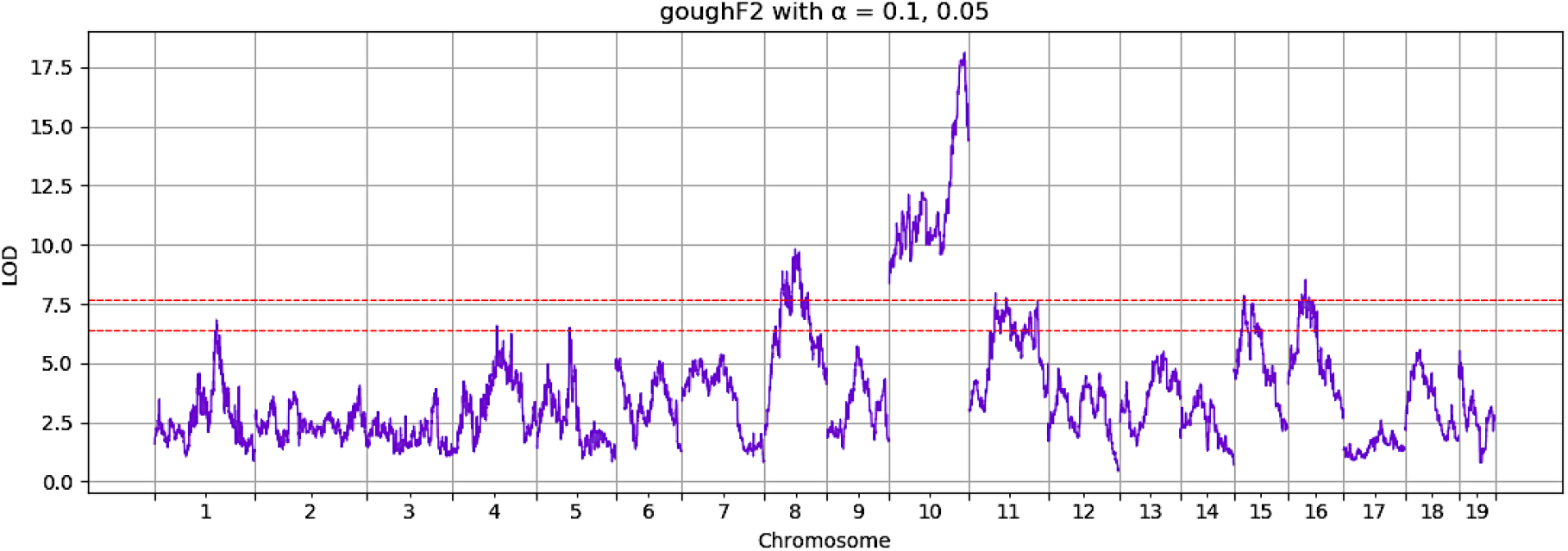
Application to the longitudinal trait data (F2 intercrosses between Gough Island mice and WSB/EiJ): LOD scores and thresholds at *α* = 0.1, 0.05

**Figure 18.**
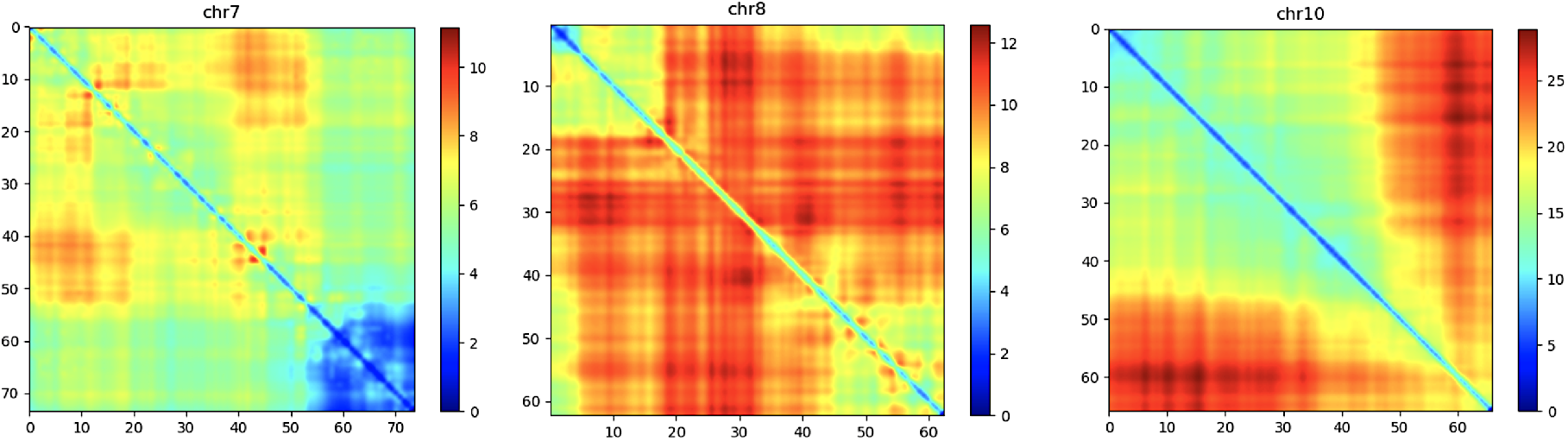
Application to the longitudinal trait data (F2 intercrosses between Gough Island mice and WSB/EiJ): LOD scores by 2D genome scan

**Figure 19.**
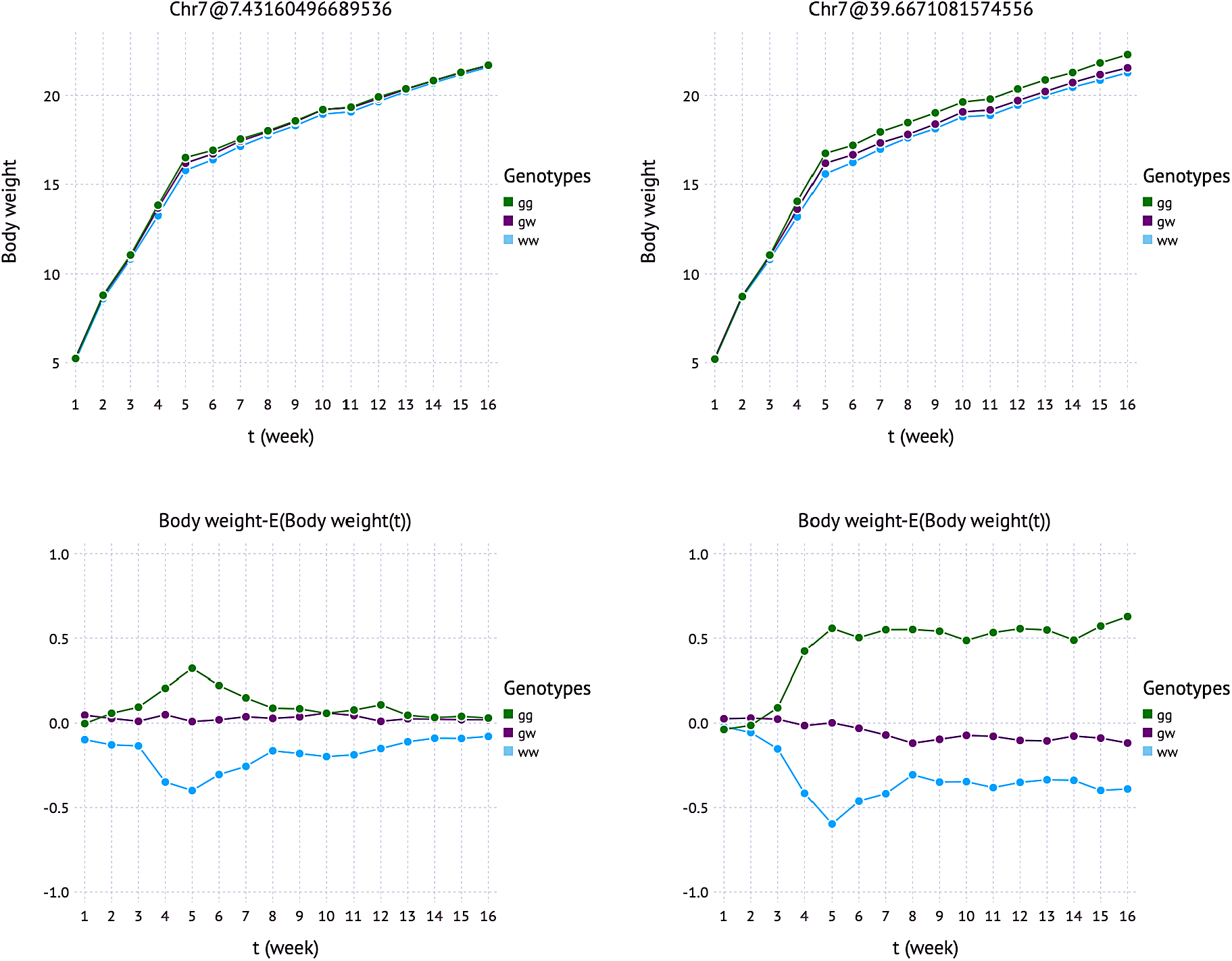
Application to the longitudinal trait data (F2 intercrosses between Gough Island mice and WSB/EiJ): Body weight effects by genotypes and corresponding deviations from weekly overall mean in Chromosome 7.

**Figure 20.**
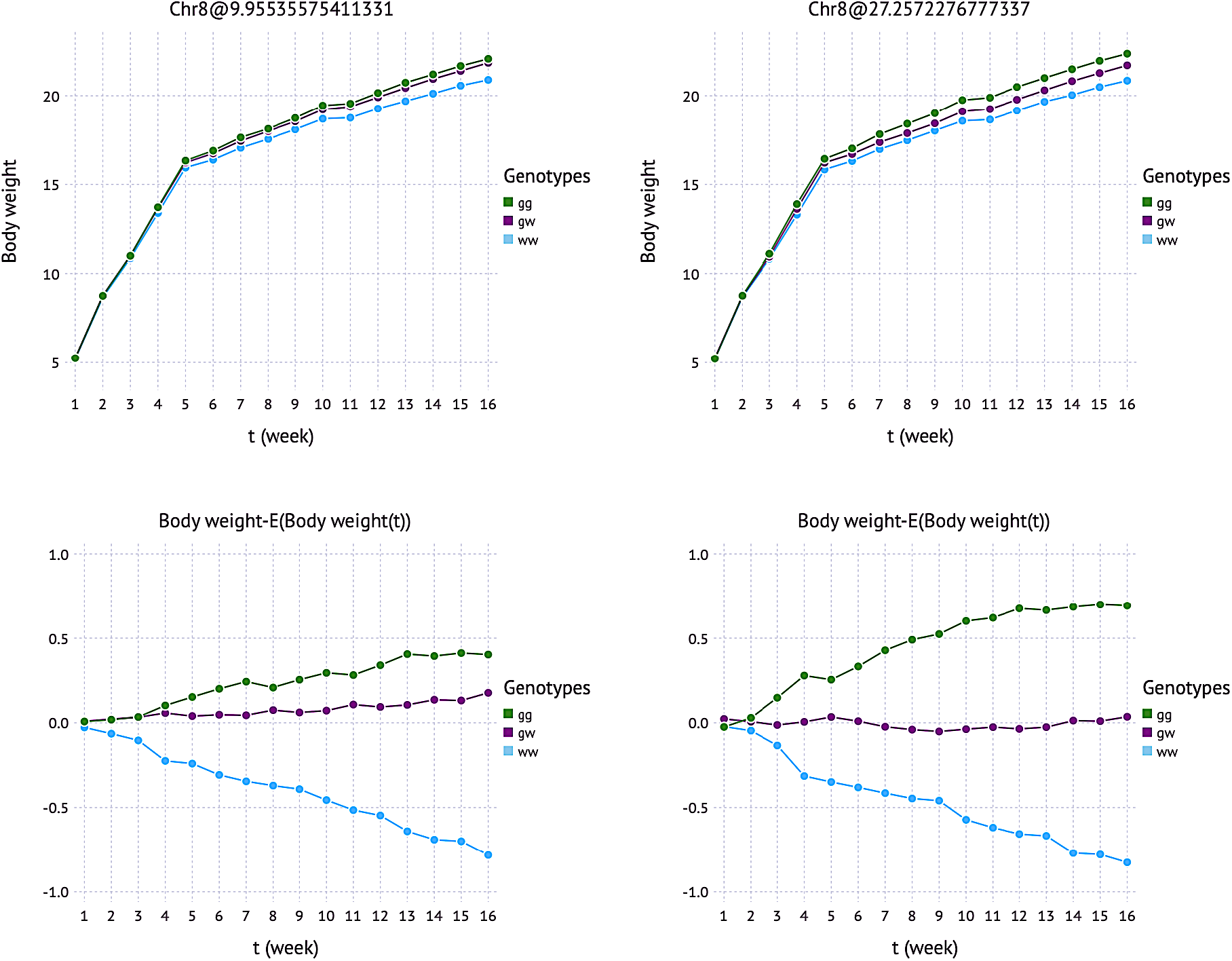
Application to the longitudinal trait data (F2 intercrosses between Gough Island mice and WSB/EiJ): Body weight effects by genotypes and corresponding deviations from weekly overall mean in Chromosome 8.

**Figure 21.**
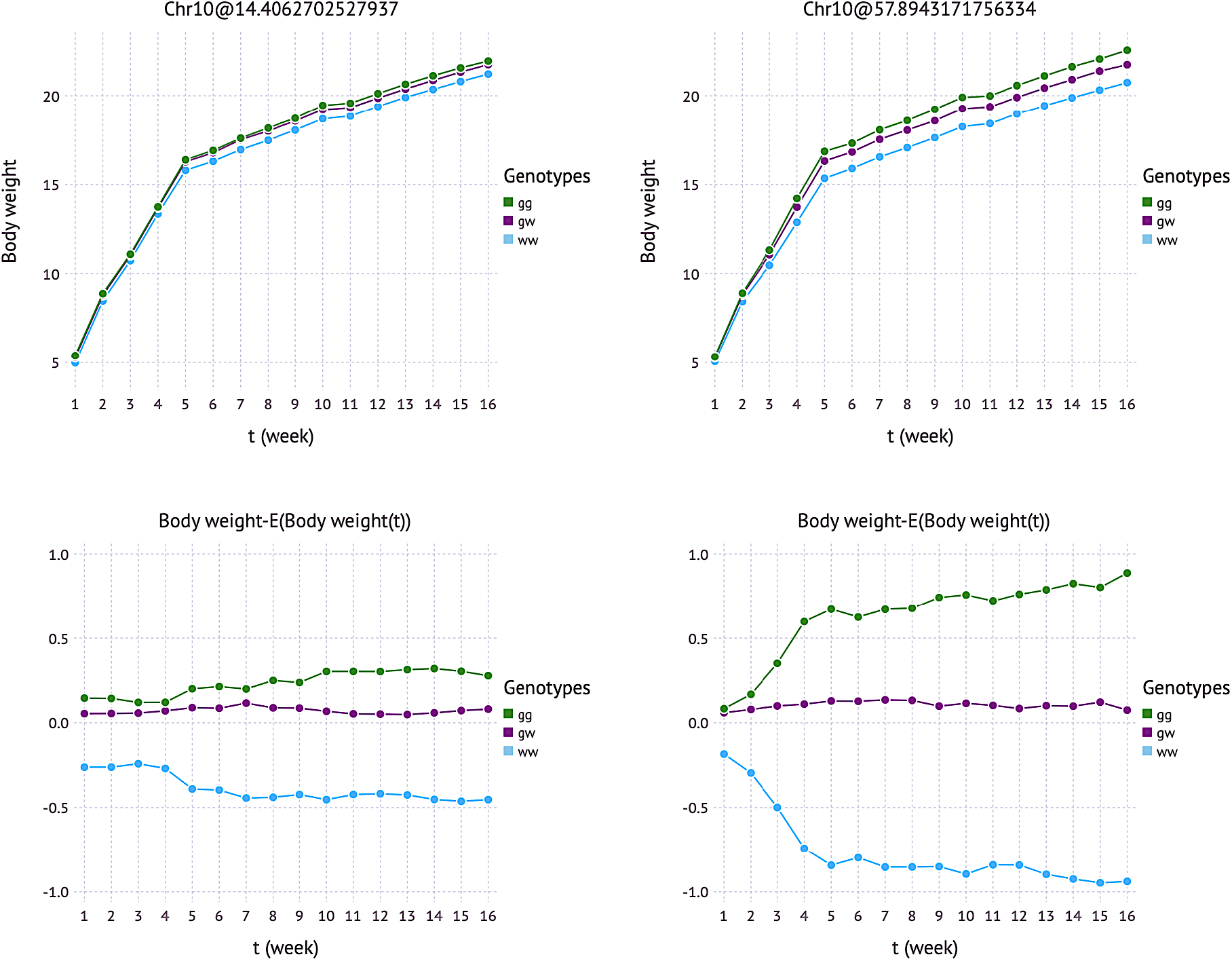
Application to the longitudinal trait data (F2 intercrosses between Gough Island mice and WSB/EiJ): Body weight effects by genotypes and corresponding deviations from weekly overall mean in Chromosome 10.

## 2 Supplementary Tables

**Table 1:**
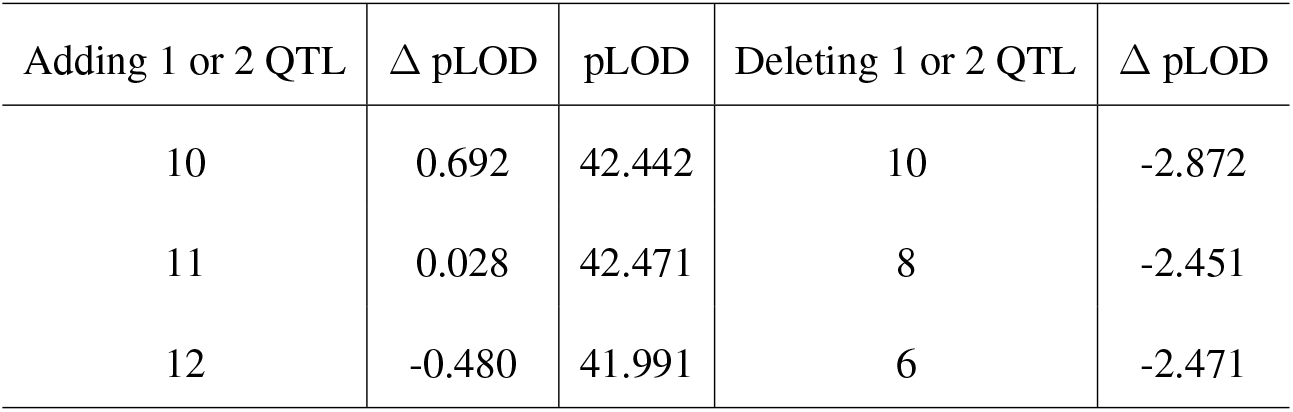
QTL selection for Arabidopsis thaliana (multiple QTL analysis) as in Supplementary Figure 15. 11 significant QTL were finalized since the largest decrease (Δ pLOD = -2.872) in the backward elimination step when dropping one QTL from 11 QTL in the forward selection step occurred.

| Adding 1 or 2 QTL | $\Delta$ pLOD | pLOD | Deleting 1 or 2 QTL | $\Delta$ pLOD |
| --- | --- | --- | --- | --- |
| 10 | 0.692 | 42.442 | 10 | -2.872 |
| 11 | 0.028 | 42.471 | 8 | -2.451 |
| 12 | -0.480 | 41.991 | 6 | -2.471 |

**Table 2:** Significant QTL for Arabidopsis thaliana as described in Supplementary Figure 15 and Supplementary Table 1.

| Chromosome | no. of QTL | position (cM) |
| --- | --- | --- |
| 1 | 2 | 57, 77 |
| 2 | 2 | 133, 140 |
| 3 | 2 | 152, 199 |
| 4 | 2 | 259, 272 |
| 5 | 3 | 280,333,353 |

## 3 Supplementary Note: Mulitivariate Linear Mixed Model for FlxQTL

The *Flex*ible multivariate linear mixed model for *QTL* mapping (FlxQTL) extended from a standard multivariate linear mixed model (MLMM) ^28^ is defined as

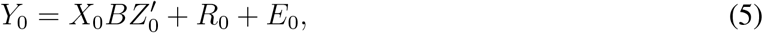

or its vectorized form,

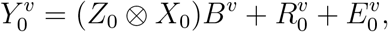

Where

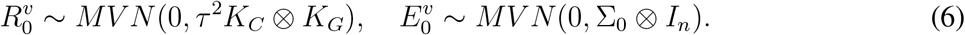

*Y*_0_ is a *n* × *m* matrix of multivariate responses (multiple traits or a trait in multiple environments), where *n* is the number of individuals (lines), *m* is the number of environments (or time points). *X*_0_ is a *n* × *p* matrix of genotypes including the intercept (or genotype probabilities) for a marker to be tested and optionally contains individual level covariates such as sex and age. *Z*_0_ is a *m* × *q* matrix of *q* trait covariates such as (site) contrasts, (orthonomal) basis functions and so on. *B*_*p*×*q*_ is then fixed effects to be estimated. *R*_0_ is a matrix of genetic random effects, and *E*_0_ is a matrix of residual errors. *K*_*G*_ is a *n* × *n* genetic relatedness (kinship) matrix, and *K*_*C*_ is a *m* × *m* trait kernel computed by information associated with the traits. In multi-environment trials (MET), the trait kernel can be a climatic relatedness matrix using high-dimensional environment information such as daily minimum or maximum temperature, precipitation, etc. *τ* ^2^ is a scalar by parameterizing a *m* × *m* convariance matrix proportional to *K*_*C*_ for dimension reduction. Note that we confine *K*_*C*_ and *K*_*G*_ to full-rank and symmetric positive definite matrices. If they are semi-positive definite, one can take advantage of a Shrinkage approach ^29, 30^ to force it to be positive definite. The eigen-decomposition to the matrices yields 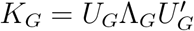 and 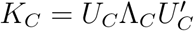. Multiplying 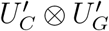 to the model gives

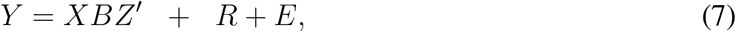

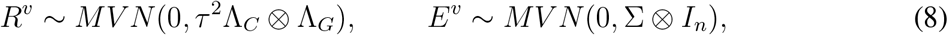

where 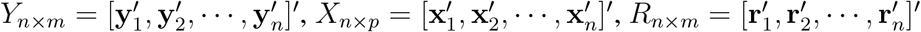, and 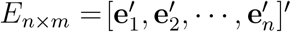.

Or, this is simplified as a vectored form,

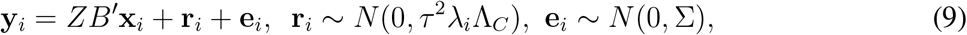

and the variance is *V*_*i*_ = *τ* ^2^*λ*_*i*_Λ_*C*_ + Σ (*i* = 1, …, *n*).

### Verification of multivariate random interaction

Assume that a multivariate random interaction term has a form of *R*_0_ = *GB*_*rand*_*C*^*′*^, where *C*_*m*×*q*1_ is *q*_1_ background (or high-dimensional) trait (or environment) information, *G*_*n*×*p*1_is *p*_1_ background genetic markers such as genotypes and genotype probabilities, and 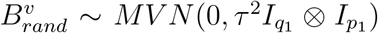. Then, the distribution of the multivariate random interactions in (6) is obtained as follows.

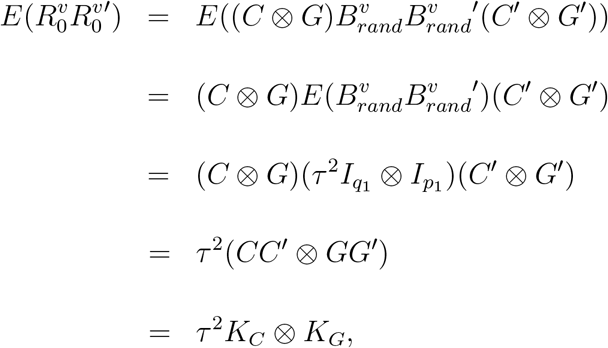

where C and G are scaled by 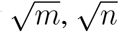, respectively.

## 4 Supplementary Note: Expectation Conditional Maximization (ECM)

Since *R* is unobservable, the joint log-likelihood function for the ECM ^14, 21^ is written as follows.

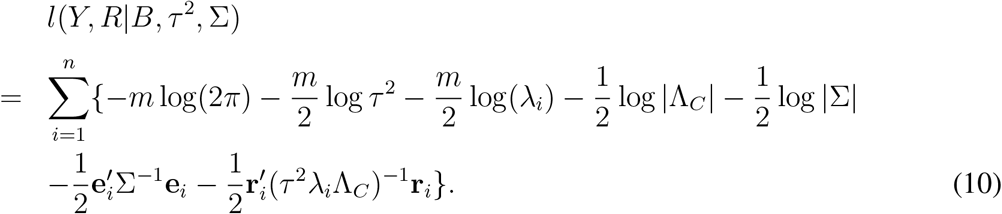

In the expectation step, the conditional distribution of **r**_*i*_ given **y**_*i*_ (*i* = 1, …, *n*), *B*^(*t*)^,*τ* ^2(*t*)^, and Σ^(*t*)^ at the current iteration *t* is ^28^

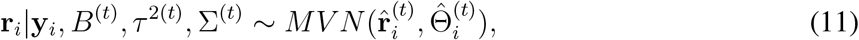

Where

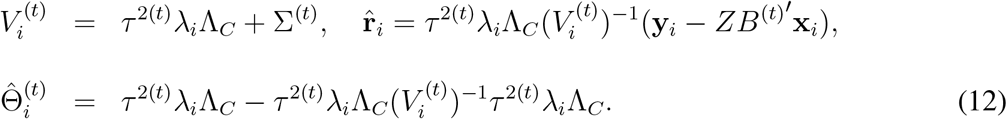

The expectation of the loglikelihood function in terms of *R* given *Y, B*^(*t*)^,*τ* ^2(*t*)^, and Σ^(*t*)^ is obtained as follows.

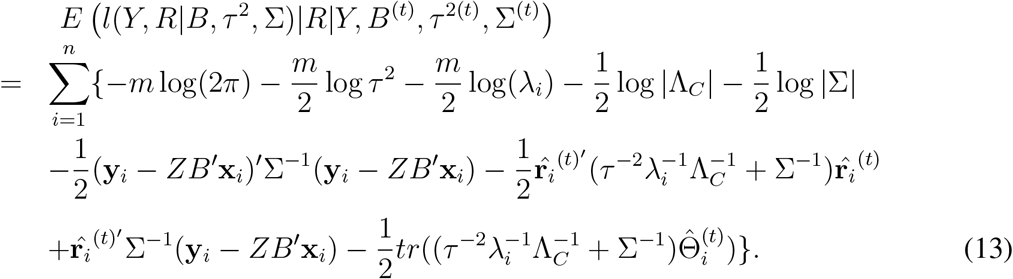

In the conditional maximization step, one can update *B*^(*t*+1)^ conditional on *τ* ^2(*t*)^ and Σ^(*t*)^, followed by updating {*τ* ^2(*t*+1)^, Σ^(*t*+1)^} conditional on *B*^(*t*+1)^, *τ* ^2(*t*)^, and Σ^(*t*) 14^.

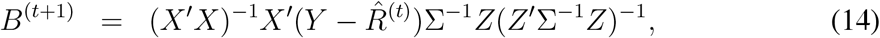

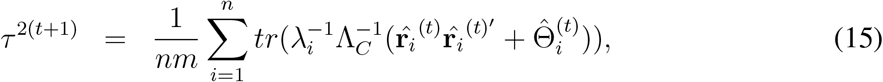

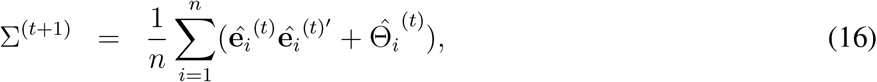

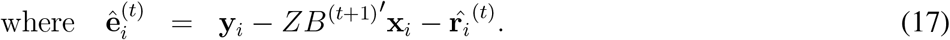

### Log-likelihood function

The joint log-likelihood function for the ECM is simplified in the following way.

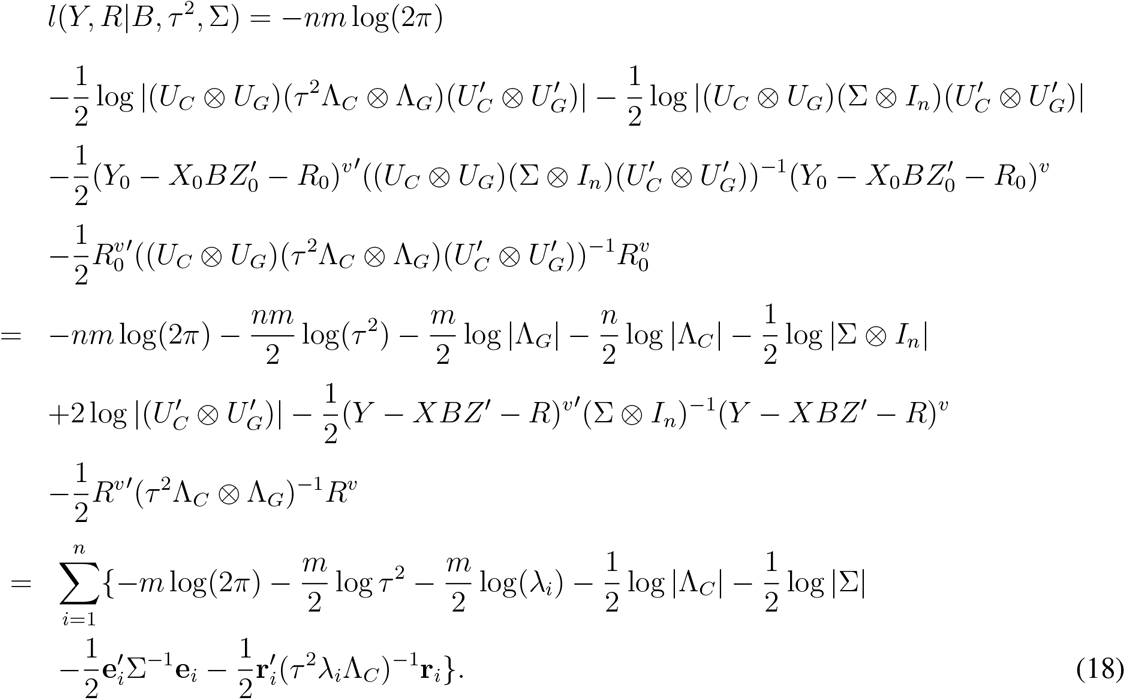

### Derivation of conditional expected log-likelihood and the closed form solutions

To obtain the closed-form solutions of *B*^(*t*+1)^, *τ* ^2(*t*+1)^, and Σ^(*t*+1)^ to the conditional expected loglikelihood in (13), one differentiates (13) with respect to each parameter after integrating out **r**_*i*_ (*i* = 1, … *n*) in the joint log-likelihood function, (10). In (10), taking expectation with respect to **r**_*i*_ (*i* = 1, …, *n*) yields

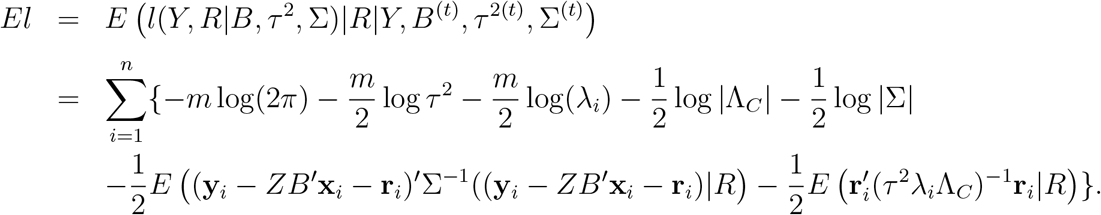

For simplicity, let *E*(·|*R*|*Y, B*^(*t*)^, *τ* ^2(*t*)^, Σ^(*t*)^) = *E*(·|*R*). The last two terms then can be simplified as follows:

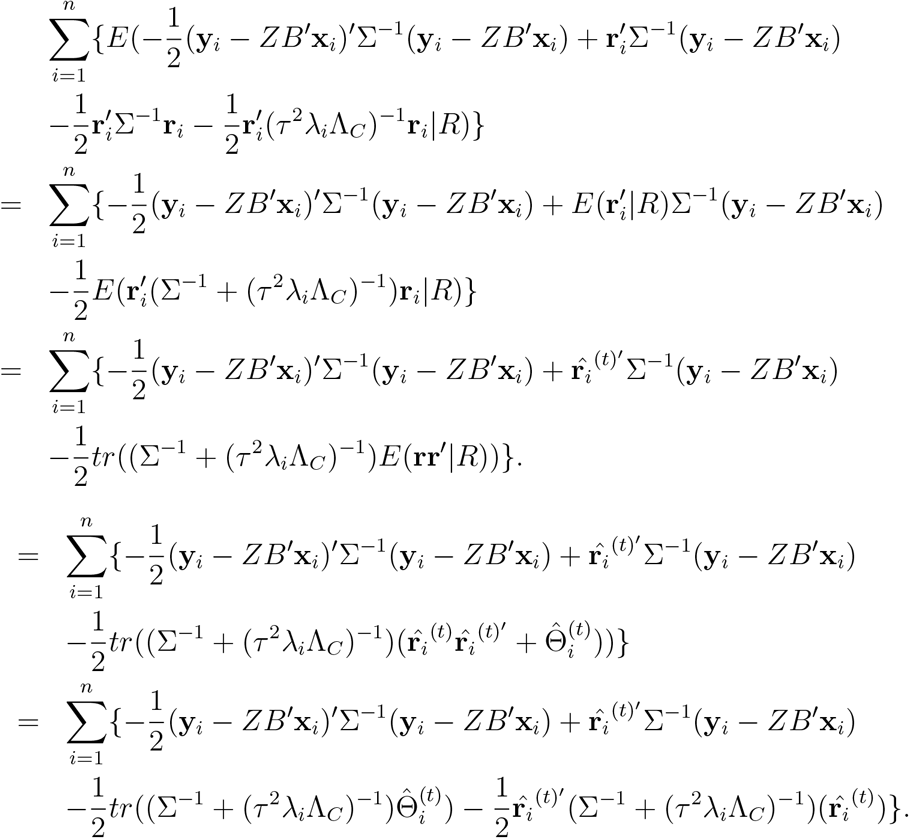

Now one can find the closed-form solutions of *B, τ* ^2^, and Σ to the expected log-likelihood function using properties of the trace, calculus for matrices. For the solution for *B*,

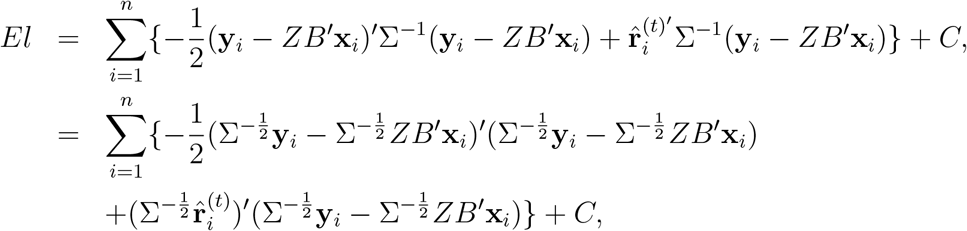

where *C* is a constant.

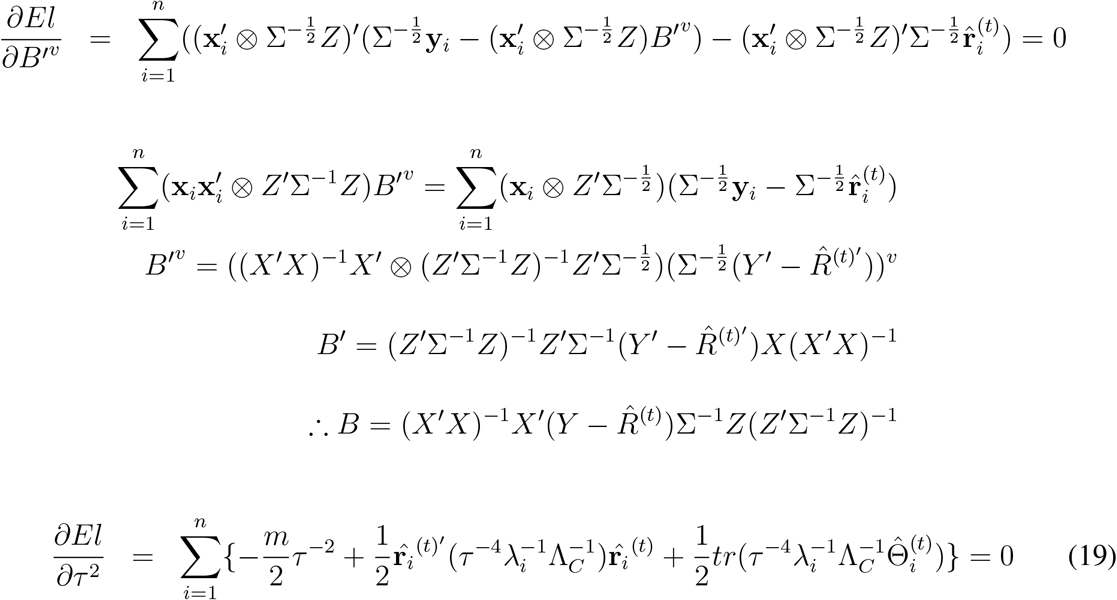

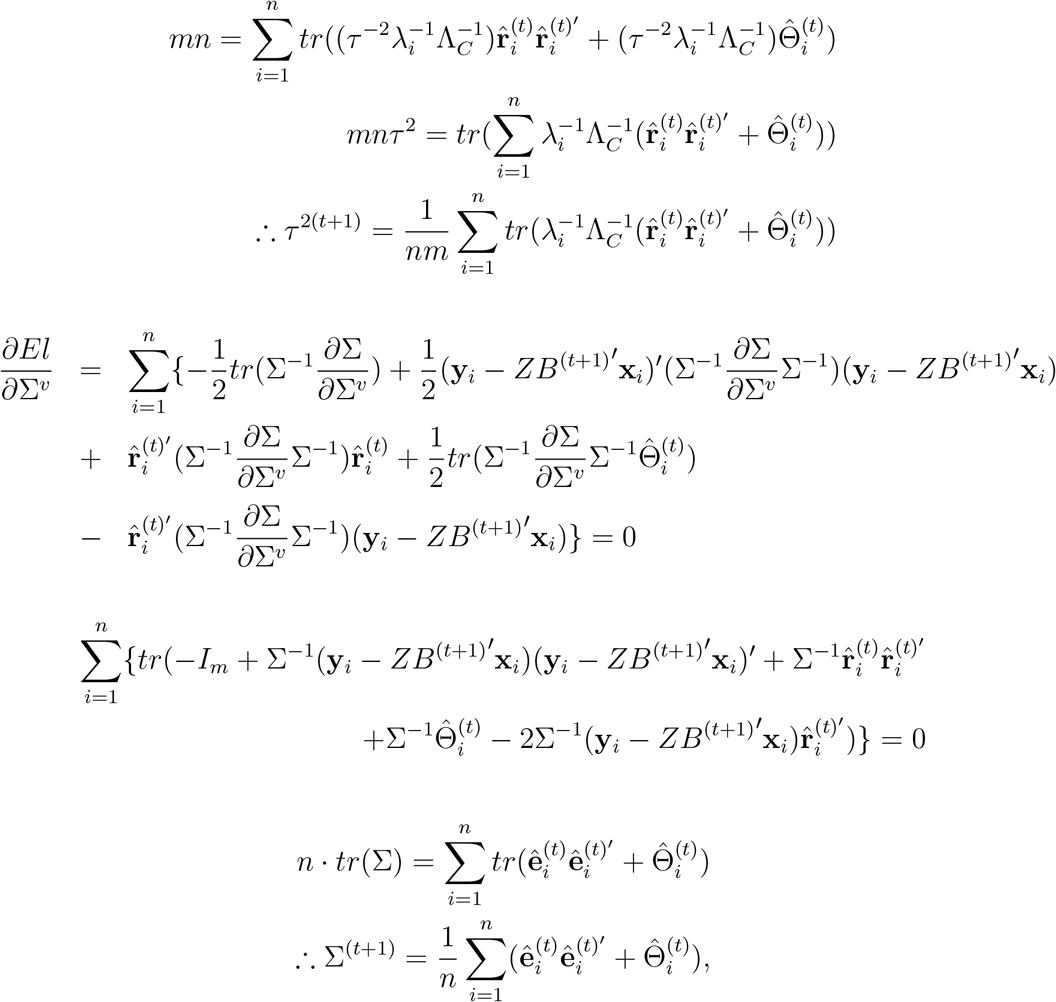

where 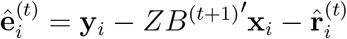.

## 5 Supplementary Note: Nesterov’s Accelerated Gradient with Speed Restart

**Algorithm**: ECM embedded in the speed restarting Nesterov’s scheme ^20^

**input**: *B*^(0)^ ∈ ℝ ^*q*×*p*^, *τ* ^2(0)^ ∈ ℝ, Σ ^(0)^ ∈ ℝ ^*m*×*m*^, *B*^(−1)^ = *B*^(0)^, *τ* ^2(−1)^ = *τ* ^2(0)^, Σ ^(−1)^ = Σ ^(0)^, *t*_*min*_ ∈ ℕ

*j* ← 1

**while** *t* = 1, | | *B*^(*t*)^ − *B*^(*t*−1)^ | | + *τ* ^2(*t*)^ − *τ* ^2(*t*−1)^ | | + | | Σ^(*t*)^ − Σ^(*t*−1)^ *> ϵ* for small enough *ϵ*

**E-step**: Integrate out the unknown complete data loglikelihood by

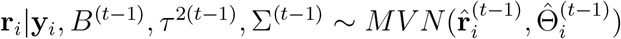 as in (11).

**CM-step**: Update each parameter by maximization over the full parameter space as in (14), (15), and (16).

**Nesterov’s Acceleration**

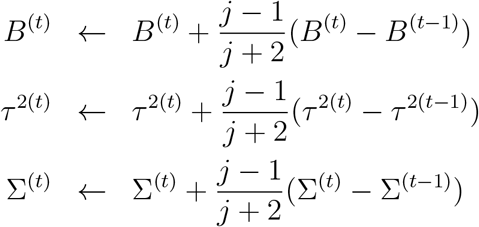

**if** | | *B*^(*t*)^− *B*^(t−1)^ | | + *τ* ^2(*t*)^− *τ* ^2(t−1)^ | | + | | Σ^(*t*)^ − Σ ^(t−1)^ | | < | | *B*^(t−1)^ − *B*^(t−2)^ | | + | | *τ* ^2(t−1)^ − *τ* ^2(t−2)^) | | + | | Σ ^(t−1)^ − Σ ^(t−2)^ | |& *t* ≥ *t*_*min*_

**then** *j* ← 1

**else** *j* ← *j* + 1

**end if**

**end while**

